# Effector-centred proximity-dependent labelling enables the discovery of cell-surface immune receptors in plants

**DOI:** 10.64898/2026.03.23.713615

**Authors:** Nicat Cebrailoglu, Esranur Budak, Sergio Landeo Villanueva, Christiaan R. Schol, Casper ter Waarbeek, Kezia Evertsz, Sjef Boeren, Matthieu H.A.J. Joosten

## Abstract

Identifying plant disease resistance proteins remains challenging, particularly for cell-surface receptors that perceive apoplastic pathogen effectors through transient or indirect interactions. Here, we establish effector-centred proximity-dependent labelling using TurboID as a protein-level approach to identify immune receptors and guarded host targets *in planta*.

We fused three apoplastic effectors, Avr2 and Avr4 from *Fulvia fulva* and XEG1 from *Phytophthora sojae*, to TurboID and transiently expressed these fusion proteins in leaves of *Nicotiana benthamiana* and tomato (*Solanum lycopersicum*). The fusions significantly biotinylate their matching receptors, including Cf-4 by Avr4-TurboID and Cf-2 by Avr2-TurboID, as well as the guarded virulence target of Avr2, the Rcr3 protease, demonstrating that both direct and indirect recognition systems can be captured.

Application of this approach to XEG1 revealed *Sl*Eix1 as the functional ortholog of NbRXEG1. Functional assays and structural modelling support *Sl*Eix1 as a signalling-competent receptor contributing to XEG1-triggered immunity. Together with previous studies, these findings position the tomato *EIX* locus as a multi-effector immune hub encoding closely related receptor-like proteins with distinct ligand specificities.

Collectively, this study establishes effector-TurboID-mediated proximity-dependent labelling as a versatile approach for identifying cell surface plant immune receptors and apoplastic virulence targets, providing a scalable route to accelerate resistance gene discovery in crop species.

## INTRODUCTION

Over the course of domestication, most crop plants have been derived from wild ancestors that evolved under strong selective pressure from diverse biotic and abiotic stresses. During modern breeding, however, selection has focused primarily on agronomically desirable traits such as yield, quality, and uniformity. Consequently, genetic diversity at loci involved in disease resistance has often been reduced or lost from cultivated gene pools. This erosion of resistance has contributed substantially to global yield losses caused by plant diseases and pests (Savary et al., 2019). Historically, these losses were mitigated largely using agrochemicals (Gruber, 2017), but increasing concerns about their environmental impact and human health risks have led to progressively stricter regulations (Donley, 2019). In the context of climate change, a growing global population, and reduced chemical inputs, durable genetic resistance has therefore become a central pillar of sustainable crop protection (Deng et al., 2020; Pathania et al., 2021). This has created an urgent need to accelerate the discovery and deployment of effective resistance (*R*) genes in crop species.

From the 1970s onwards, wild relatives of crop plants have been extensively exploited as sources of disease resistance, which were introgressed into elite cultivars through crossing (Schouten et al., 2019; Chauhan et al., 2019; Gramazio et al., 2021). Initially, resistance was tracked mainly by phenotypic disease assays, while later PCR-based markers were used to track resistance-associated chromosomal regions. Although marker-assisted selection enabled the transfer of *R* loci across generations, such loci are often large and may contain multiple candidate genes. Moreover, recombination events or mutations can uncouple markers from the causal gene, resulting in the loss of resistance while marker analyses remain positive (Collard & Mackill, 2008). Consequently, cloning and functional characterisation of the underlying *R* gene have become essential for robust resistance breeding.

Plant immunity is commonly described as a two-layered system. In the first layer, cell-surface pattern-recognition receptors (PRRs) detect conserved microbe- or damage-associated molecular patterns (MAMPs/DAMPs), thereby activating pattern-triggered immunity (PTI) (Jones & Dangl, 2006; Bigeard et al., 2015; Han, 2019). In the second layer, immune responses are often mediated by intracellular nucleotide-binding leucine-rich repeat (NLR) proteins that monitor cytoplasmic pathogen effectors or their perturbation of host targets, leading to effector-triggered immunity (ETI). Although ETI is frequently associated with NLRs, cell-surface receptors can also contribute to effector recognition (Jones & Dangl, 2006; van der Burgh & Joosten, 2019; Ngou et al., 2022). According to the classical gene-for-gene model, resistance is activated when a plant R protein recognises a matching pathogen avirulence (Avr) protein (Flor, 1971). Many Avr proteins are secreted effectors deployed by pathogens to suppress host immunity and facilitate colonisation. These effectors act either in the apoplast or within the host cells, and in their turn, plants have evolved surveillance systems to perceive the presence or activity as a virulence factor of these effectors (Jones & Dangl, 2006; Dodds & Rathjen, 2010).

Plasma membrane (PM)–localised receptor-like proteins (RLPs) and receptor-like kinases (RLKs) are responsible for perceiving apoplastic effectors, whereas cytoplasmic effectors are typically recognised by NLRs. Recognition of an effector imposes strong selective pressure on pathogen populations, favouring races that evade detection through loss or modification of the recognised effector, a process known as resistance breakdown (Joosten et al., 1994; McDonald & Linde, 2002).

Early effector discovery relied on comparative analyses of virulent and avirulent pathogen races, leading to the identification of numerous bacterial and fungal effectors (Napoli & Staskawicz, 1987; van Kan et al., 1991; Luderer & Joosten, 2001). With the advent of high-throughput sequencing and advanced proteomics, entire pathogen effectoromes can now be predicted and screened *in planta* using effectoromics approaches (Mesarich et al., 2018). However, despite rapid advances in effector discovery, identifying the corresponding plant immune receptors remains challenging, as classical genetic mapping of *R* loci is time-consuming and poorly suited for systematic effector–receptor matching (Kourelis & van der Hoorn, 2018; Chitwood-Brown et al., 2021).

Several sequence-based approaches have markedly accelerated *R* gene cloning. *R* gene enrichment sequencing (RenSeq) enables targeted capture of NLR- and RLP/RLK-encoding loci to improve genome annotations and facilitate rapid mapping (Jupe et al., 2013; Andolfo & Ercolano, 2015; Chen et al., 2018; Lin et al., 2020). Mutagenesis-coupled RenSeq (MutRenSeq) combines exome capture with chemical mutagenesis and has enabled rapid cloning of multiple rust and mildew *R* genes in wheat (Steuernagel et al., 2016; Zhang et al., 2020). Association genetics RenSeq (AgRenSeq) integrates *k*-mer–based association mapping with RenSeq to clone *NLRs* directly from diverse germplasm panels (Arora et al., 2019). While these approaches have transformed *R* gene discovery, they primarily operate at the DNA level and depend on complex informative sequence variation, rather than directly interrogating the protein complexes that assemble around pathogen effectors.

Because classical genetics and sequence-based methods alone are insufficient to rapidly match effectors to their corresponding receptors, proteomic approaches are increasingly being explored. Affinity purification methods, such as GREEN FLUORESCENT PROTEIN (GFP) pull-downs, have been widely used to identify interacting proteins. However, effector–receptor interactions are not always direct, and the interaction can also be weak and transient, resulting in loss of the interaction upon a regular pull-down (Göös et al., 2022). For example, the apoplastic protease REQUIRED FOR CLADOSPORIUM RESISTANCE 3 (Rcr3) is inhibited by the *F. fulva* effector Avr2, and this perturbation is monitored by the R protein Cf-2 (Dixon et al., 1996), which does not bind Avr2 directly (Luderer et al., 2002; Rooney et al., 2005). Such guard or decoy mechanisms illustrate that R proteins often sense effector-induced modifications of host targets rather than the effector itself (Kourelis & van der Hoorn, 2018). Consequently, methods that rely on stable, direct interactions may fail to capture key components of immune recognition.

Proximity-dependent labelling (PL) offers a powerful alternative for mapping local protein environments *in vivo*. In PL approaches, a promiscuous biotin ligase is fused to a bait protein, enabling covalent biotinylation of nearby proteins within a limited radius. These biotinylated proteins can then be enriched and identified by mass spectrometry (Roux et al., 2012). Early PL enzymes such as BioID (Lin et al., 2017), were of limited utility in plants, but the engineered biotin ligase TurboID displays high catalytic activity with low toxicity and has proven suitable for transient expression in plant tissues (Branon et al., 2018; Zhang et al., 2019). In recent studies, TurboID has been fused to pathogen effectors to identify host susceptibility factors and downstream signalling components (Shi et al., 2023; Hawk et al., 2023). However, effector–TurboID fusions have not yet been systematically exploited to identify the immune receptors that perceive apoplastic effectors or guard their virulence targets in crop plants. An approach that operates directly at the protein level and captures both direct and indirect interactions within effector-associated complexes would provide a valuable complement to existing sequence-based *R* gene discovery strategies.

Here, we employ effector-centred PL using TurboID to identify immune receptors in *Nicotiana benthamiana (N. benthamiana, Nb)* and tomato (*Solanum lycopersicum, Sl*). For this, we fused three apoplastic effectors—Avr2 and Avr4 from the fungal tomato pathogen *Fulvia fulva* (Joosten et al., 1994; Luderer et al., 2002) and XEG1 from the oomycete pathogen *Phytophthora sojae* (Ma et al., 2015) —to TurboID and expressed them transiently *in planta*. Avr2 and Avr4 were selected as benchmark effectors with well-characterised receptors, being Cf-2 (Dixon et al., 1996) and Cf-4 (Thomas et al., 1997), respectively, thus enabling validation of the approach for both an indirect and a presumed direct recognition system. The glycoside hydrolase family 12 member with xyloglucanase activity, XEG1, matching the known RLP RESPONSE TO XEG1 (RXEG1) of *N. benthamiana* (Wang et al., 2018) was included to address a biologically relevant case, as recognition of XEG1 by tomato has been demonstrated, but the tomato receptor had not yet been conclusively identified at the beginning of this study. By combining effector-centred PL with functional assays and structural analyses, we demonstrate that this strategy enables the identification of cell-surface immune receptors and guarded host targets, thereby providing a scalable framework to accelerate *R* gene discovery in crop species.

## RESULTS

### Effector–TurboID fusion proteins localise to the apoplast and biotinylate their matching receptors in *N. benthamiana*

To establish whether apoplastic effectors can be used as PL baits for cell-surface immune receptors, the fungal effectors Avr4 and Avr2 and the oomycete GH12 protein XEG1 were fused C-terminally to YELLOW FLUORESCENT PROTEIN (YFP) and TurboID. The resulting effector–YFP–TurboID fusion proteins were transiently expressed in *N. benthamiana* leaves by *Agrobacterium tumefaciens*–mediated transient transformation (agroinfiltration). To assess whether effector–TurboID fusion proteins accumulate in the apoplast, apoplastic fluid (AF) was isolated from leaves expressing Avr4–YFP–TurboID and was analysed by immunoblotting using an anti-TurboID antibody. A prominent band of ~80kDa, corresponding to full-length Avr4–YFP–TurboID fusion protein, was detected (Figure 1A), together with additional bands of approximately 70kDa and 36kDa (Figure 1A), consistent with YFP–TurboID and TurboID fragments, respectively. These data indicate that the effector–TurboID fusion protein accumulates in the apoplast but undergoes at least partial proteolytic processing. Consistent with this interpretation, co-expression of the *Phytophthora infestans* extracellular protease inhibitor EPI1 (Tian et al., 2004) increased the accumulation of the full-length fusion protein (Figure S1), indicating substantial apoplastic proteolysis of Avr4–YFP–TurboID.

**Figure 1.**
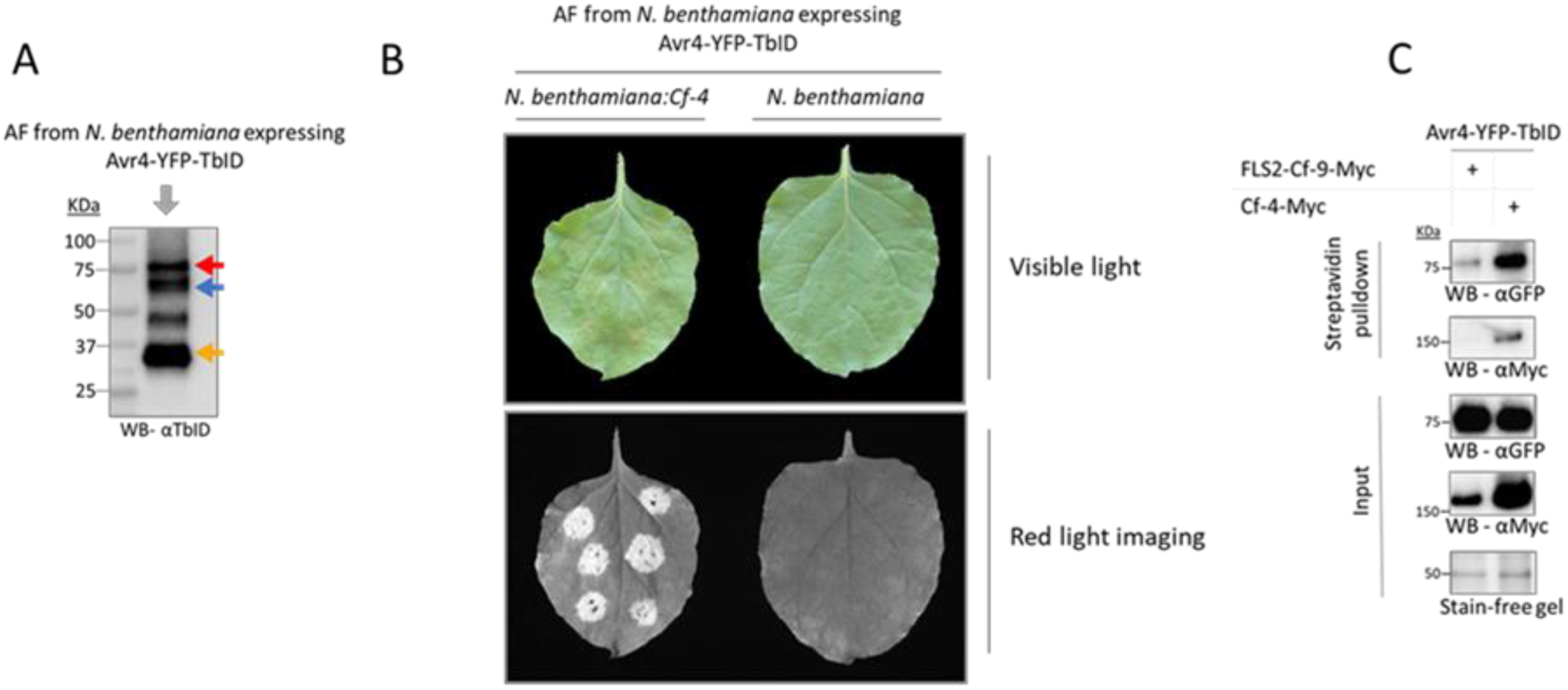
PL of the cell-surface receptor Cf-4 by apoplastic Avr4–YFP–TurboID. (A) Apoplastic fluid (AF) isolated from *N. benthamiana* leaves transiently expressing Avr4–YFP–TurboID was analysed by SDS–PAGE, followed by immunoblotting using an anti-TurboID antibody (αTbID). A band of ~80kDa corresponding to the full-length Avr4–YFP–TurboID fusion protein (red arrow) was detected, together with lower-molecular-weight bands consistent with the migration of YFP–TurboID (blue arrow) and TurboID (yellow arrow), indicating partial proteolytic processing in the apoplast. (B) AF containing Avr4–YFP–TurboID was infiltrated into WT and stable *Cf-4*–transgenic *N. benthamiana* leaves (*N. benthamiana:Cf-4*). A HR was observed exclusively in the Cf-4–expressing plants, as visualised under visible light and by red light imaging at 2 dpi, demonstrating retention of Avr4 elicitor activity. (C) Avr4–YFP–TurboID was co-expressed with Cf-4–Myc or FLS2–Cf-9–Myc in *N. benthamiana*. At 2 dpi, leaves were supplied with biotin and biotinylated proteins were enriched from total protein extracts using streptavidin-coated beads. Immunoblotting with anti-GFP confirmed the presence of Avr4–YFP–TurboID in the streptavidin-enriched fraction, while anti-Myc immunoblotting detected Cf-4–Myc, but not FLS2–Cf-9–Myc, in the pull-down, despite both receptors being present in the input. A stain-free gel is shown as a loading control. The prominent band at approximately 50kDa corresponds to Rubisco, indicating comparable protein loading of both samples. These data show that apoplastic Avr4–TurboID specifically biotinylates its matching receptor Cf-4.

To test whether apoplast-localised Avr4–YFP–TurboID retains its elicitor activity, AF containing Avr4–YFP–TurboID was infiltrated into leaves of stable transgenic *N. benthamiana* plants expressing the *Cf-4* immune receptor (*N. benthamiana:Cf-4*) (Gabriëls et al., 2006) and into wild-type (WT) plants. A hypersensitive response (HR) was observed specifically in *Cf-4*–expressing plants but not in the WT control, as visualised by red fluorescence imaging (Landeo Villanueva et al., 2021) (Figure 1B). These results demonstrate that the Avr4–YFP–TurboID fusion protein is recognised by Cf-4 when secreted into the apoplast.

Next, we tested whether Avr4–YFP–TurboID biotinylates its matching receptor Cf-4. For this, Avr4–YFP–TurboID was transiently co-expressed in *N. benthamiana* with Cf-4–Myc or with FLS2–Cf-9–Myc, which is a chimeric receptor consisting of the FLAGELLIN-SENSING 2 (FLS2) ectodomain, fused to the Cf-9 endodomain (Wu et al., 2019). FLS2–Cf-9–Myc localises to the PM and interacts with the adaptor RLK SUPPRESSOR OF BIR1-1 (SOBIR1), similar to Cf-4 (Liebrand et al., 2013; Wu et al., 2019), and therefore served as a negative control. Anti-GFP immunoblotting confirmed the presence of Avr4–YFP–TurboID in the streptavidin-enriched fraction, while anti-Myc immunoblotting detected Cf-4–Myc, but not FLS2–Cf-9–Myc, in the pull-down, despite both receptors being present in the input samples (Figure 1C). These results demonstrate that apoplastic Avr4–TurboID specifically biotinylates its matching receptor Cf-4. Importantly, the absence of FLS2–Cf-9–Myc labelling indicates that Avr4–YFP–TurboID does not promiscuously biotinylate non-matching receptors.

Proper apoplastic localisation and specific RXEG1 receptor biotinylation were also observed for XEG1–YFP–TurboID when transiently co-expressed with *Nb*RXEG1-Myc (Figures S2 and S3). Avr2–TurboID–mediated PL was not assessed in *N. benthamiana* due to the reported dysfunction of the *N. benthamiana* Rcr3 ortholog in Avr2/Cf-2–mediated immunity (Kourelis et al., 2020). Still, all three effector–TurboID fusion proteins were detected in the AF of agroinfiltrated *N. benthamiana* leaves (Figure S3).

### PL identifies endogenous receptors matching apoplastic effectors in *N. benthamiana*

To determine whether effector–TurboID fusions can identify endogenous immune receptors without prior transient receptor overexpression, Avr4–TurboID and XEG1–TurboID were transiently expressed in *N. benthamiana:Cf-4*, after which the samples were prepared for LC-MS/MS following the procedure described in the Materials and Methods section. For each effector, the analysis of the raw LC-MS/MS data revealed reproducible enrichment patterns across all biological replicates, as confirmed by principal component analysis (Figure S5). Volcano plot analysis showed that each effector–TurboID fusion strongly enriched its corresponding bait protein, consistent with efficient self-biotinylation (Figure 2). Importantly, the matching immune receptors Cf-4 and *Nb*RXEG1 were significantly enriched in the Avr4–TurboID and XEG1–TurboID datasets, respectively. Although these receptors did not form extreme outliers in the volcano plots, they were among the few receptor-like proteins present within the significantly enriched fraction and were readily identifiable based on functional annotation (Supplementary Data 1). The recovery of Cf-4 and *Nb*RXEG1 demonstrates that effector–TurboID PL detects endogenous matching receptors in *N. benthamiana*, without transient overexpression of the receptor-encoding genes. This result extends the targeted co-expression assay, where Cf-4 and RXEG1 are expressed from a 35S promoter, and highlights the ability of effector-centred PL to identify native endogenous immune receptors directly *in planta*.

**Figure 2.**
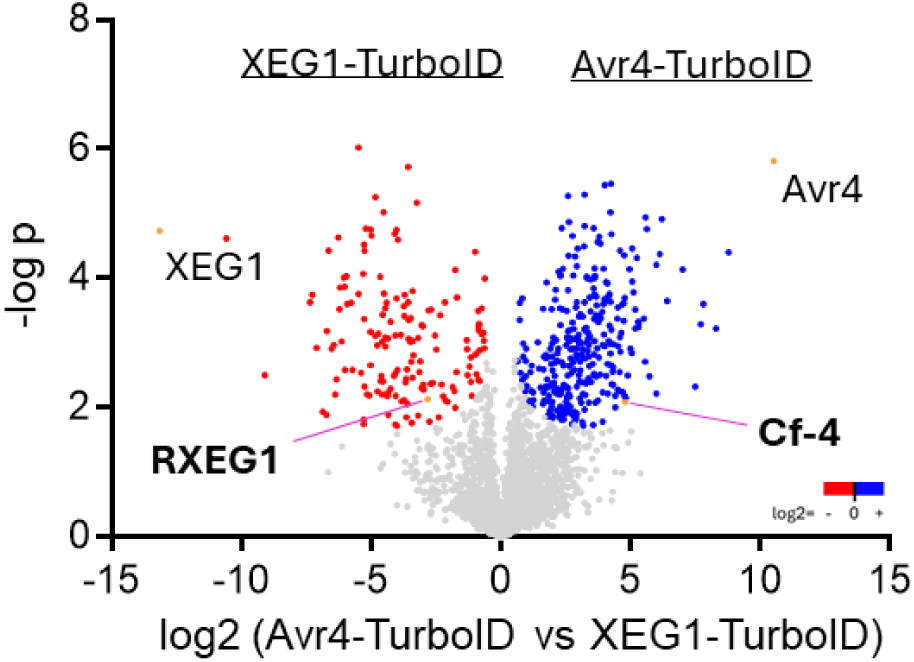
Effector–TurboID fusions biotinylate their matching endogenous receptors in *N. benthamiana*. Shown are representative volcano plots of proteins identified by PL using XEG1–TurboID (left) and Avr4–TurboID (right), transiently expressed in *N. benthamiana:Cf-4*. Log2-transformed label-free quantification (LFQ) intensity ratios are shown on the x-axis and −log10 (*p*) values on the y-axis. Proteins significantly enriched in the XEG1–TurboID- and Avr4–TurboID-generated datasets are shown in red and blue, respectively (two-sided Student’s *t*-test with permutation-based FDR < 0.05, S₀ = 0.1), while non-significantly enriched proteins are shown in grey. The TurboID-fused bait proteins are among the most strongly enriched proteins in each dataset. Notably, the known endogenous receptors *Nb*RXEG1 and Cf-4 are significantly enriched in the XEG1–TurboID- and Avr4–TurboID-generated datasets, respectively, and are readily distinguishable from the background proteins. Data originate from three independent biological replicates per effector. All identified proteins upon Perseus analysis are listed in Supplementary Data 1.

### Tomato is a suitable host for apoplastic PL

To assess whether effector-centred PL can also be applied in a crop species, the approach was extended to tomato, and Avr2 was included as a third effector. Protein expression and the appropriate subcellular localisation of the effector–TurboID fusion proteins were confirmed by confocal microscopy, which revealed robust YFP fluorescence for Avr4–, Avr2–, and XEG1–YFP-TurboID fusions (Figure S6). To validate effector–TurboID expression and proper enzymatic activity prior to PL–MS, the three fusion proteins were transiently expressed in leaves of the near-isogenic tomato lines Moneymaker (MM)–Cf-4 (MM-Cf-4) and MM–Cf-2 (MM-Cf-2) (Boukema, 1977), and total protein extracts were analysed by western blotting. Because TurboID activity requires ATP to generate a reactive biotin–AMP intermediate (Branon et al., 2018), the biotin solution that was infiltrated into the leaves before harvesting was supplemented with ATP to compensate for the low availability of ATP in the apoplast. All effector–TurboID fusions were expressed at detectable levels and showed robust *in planta* biotinylation activity (Figure 3A). Quantification of the streptavidin–HRP signals revealed that extracellular ATP consistently enhanced overall protein biotinylation across all effector–TurboID samples (Figure 3B), demonstrating that ATP supplementation indeed improves PL efficiency in tomato.

**Figure 3.**
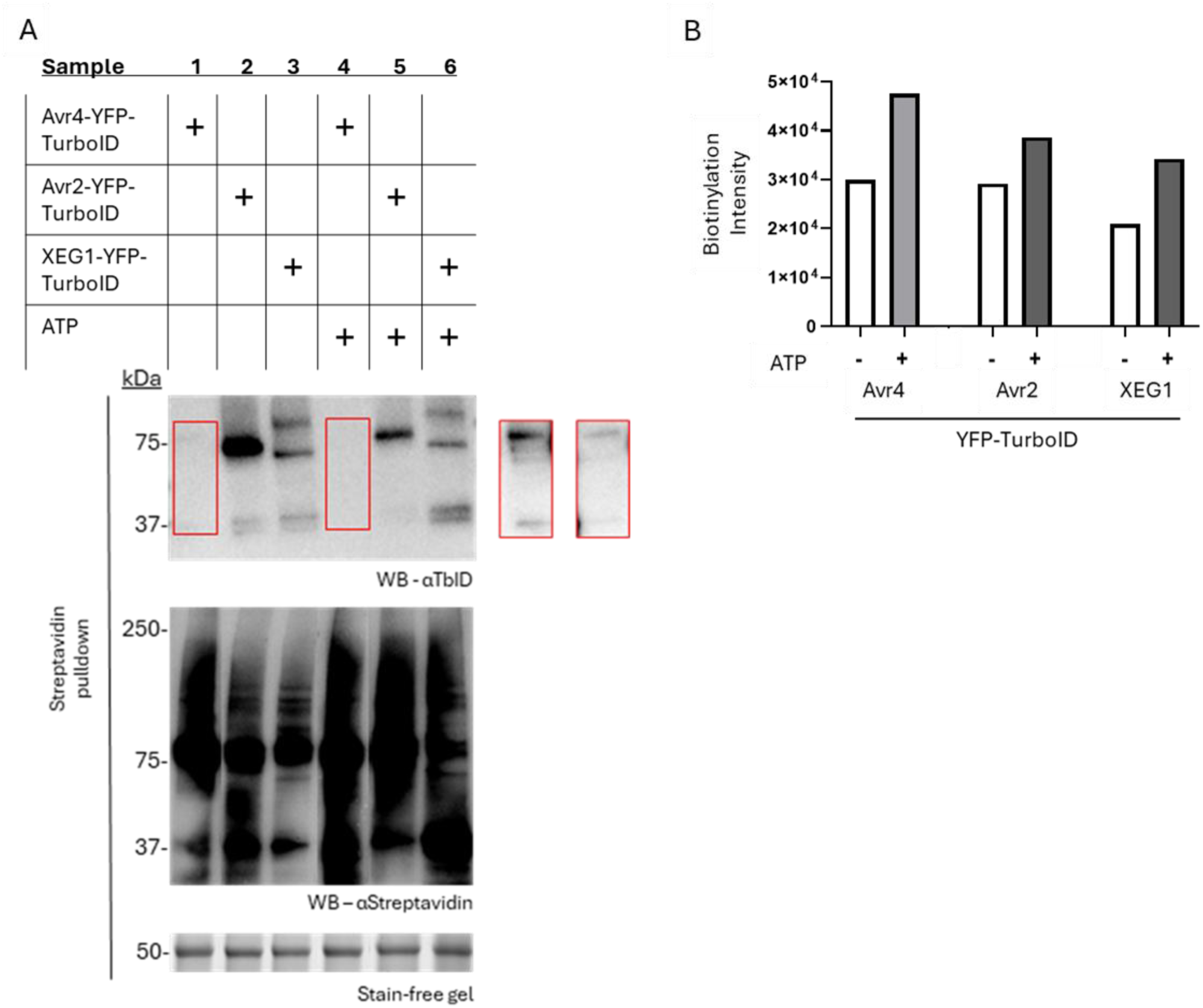
Expression and biotinylation activity of effector–TurboID fusion proteins in tomato. Effector–TurboID fusion proteins were transiently expressed in tomato leaves by agroinfiltration. All leaves were infiltrated with biotin prior to harvest; where indicated, leaf tissue was additionally treated with extracellular ATP. Plus (+) symbols indicate expression of the indicated effector–YFP–TurboID construct. (A) Total protein extracts were analysed by western blotting using αTbID antibodies to confirm expression of the effector–TurboID fusion proteins (upper panel) and using streptavidin–HRP to visualise biotinylated proteins (middle panel). Red boxes indicate Avr4–YFP–TurboID bands that were only weakly or not visible upon automatic exposure, but detectable upon longer exposure. Lanes corresponding to samples 1 and 4 of the streptavidin blot were rearranged to reflect the correct sample order; all lanes originate from the same gel and exposure. In the lower panel, the prominent band at approximately 50kDa corresponds to Rubisco and serves as a loading control. (B) Overall biotinylation activity was quantified by densitometric analysis of the streptavidin–HRP signal intensities shown in the middle panel of (A). Note that extracellular ATP supplementation consistently increased the overall biotinylation levels across all effector–TurboID-generated samples, indicating enhanced TurboID-mediated PL in tomato tissue.

When transiently expressed in MM–Cf-4 leaves, all effector–TurboID fusions exhibited strong self-enrichment in the volcano plot of the LC-MS/MS data (Figure 4). Furthermore, Avr4–TurboID distinctly enriched the receptor Cf-4 among the significantly enriched proteins, whereas Avr2–TurboID enriched its known virulence target Rcr3 and additional apoplastic proteases (Supplementary Data 1), consistent with Avr2 functioning as a cysteine protease inhibitor and the established interaction of Avr2 with Rcr3 (Rooney et al., 2005; van Esse et al., 2008; Shabab et al., 2008). Compared with *N. benthamiana:Cf-4*, enrichment of Cf-4 in tomato was more pronounced, coinciding with ATP supplementation and suggesting an improved labelling efficiency under these conditions.

**Figure 4.**
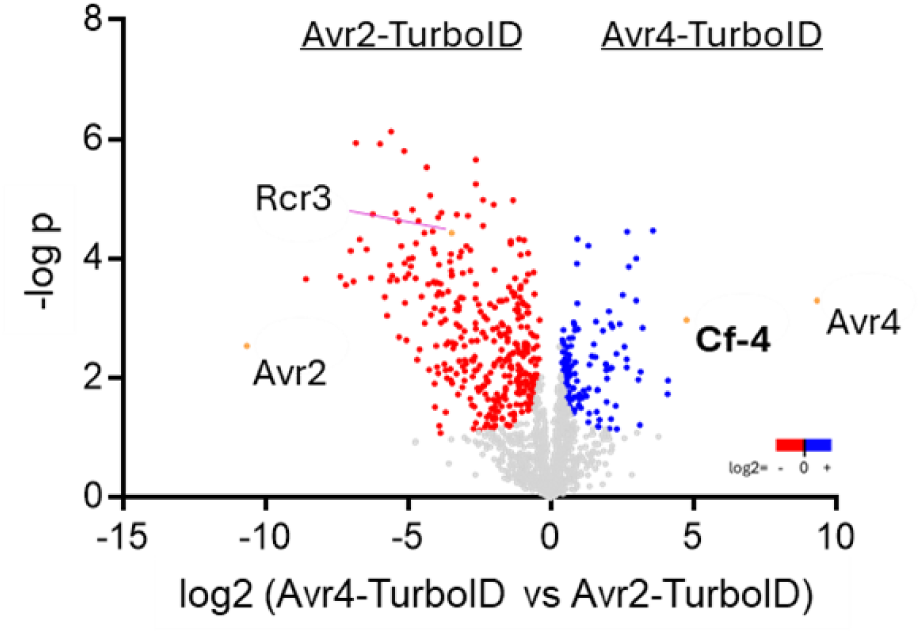
Effector–TurboID fusions biotinylate matching receptors and virulence targets in MM–Cf-4 tomato plants. Volcano plots of PL–MS data obtained following transient expression of Avr2–TurboID (left) or Avr4–TurboID (right) in leaves of MM–Cf-4 tomato plants. Log2-transformed LFQ intensity ratios are plotted against −log10 (*p*) values. Proteins significantly enriched in the Avr2–TurboID- and in the Avr4–TurboID-generated samples are shown in red and blue, respectively (two-sided Student’s *t*-test with permutation-based FDR < 0.05, S₀ = 0.1). Cf-4 is distinctly enriched in the Avr4–TurboID-generated samples, whereas Rcr3 and additional proteases are enriched in Avr2–TurboID-generated samples. The data represent three biological replicates, each prepared from five infiltrated leaves.

When expressed in MM–Cf-2 leaves, Avr2–TurboID again showed efficient self-enrichment and also significantly enriched Cf-2 (Figure 5A). Although Cf-2 did not emerge as a strong outlier in the volcano plot, it was among the few RLPs in the significantly enriched fraction and could be readily identified based on annotation. In contrast, Rcr3 was not significantly enriched under these conditions. Avr4–TurboID, included as a negative control in the Cf-2 background, did not enrich neither Cf-2 nor Rcr3, demonstrating effector–receptor specificity. In the MM–Cf-2 background, although some RLPs were significantly enriched, none of them was similar to *Nb*RXEG1. However, a detailed inspection of the complete dataset revealed that the only RLP that has similarity to *Nb*RXEG1 (Q6JN47) was detected exclusively in the XEG1–TurboID samples (Figure 5B). Subsequent annotation identified this protein as the tomato ETHYLENE-INDUCING XYLANASE receptor *Sl*Eix1 (Solyc07g008620), previously proposed to function as a decoy receptor (Ron & Avni, 2004). Peptides corresponding to *Sl*Eix1 were only detected in all three XEG1–TurboID biological replicates (Table S1) but failed to pass the statistical significance threshold due to low (<3) peptide counts in one replicate. Together, these data identify *Sl*Eix1 as a candidate XEG1-associated RLP in tomato.

**Figure 5.**
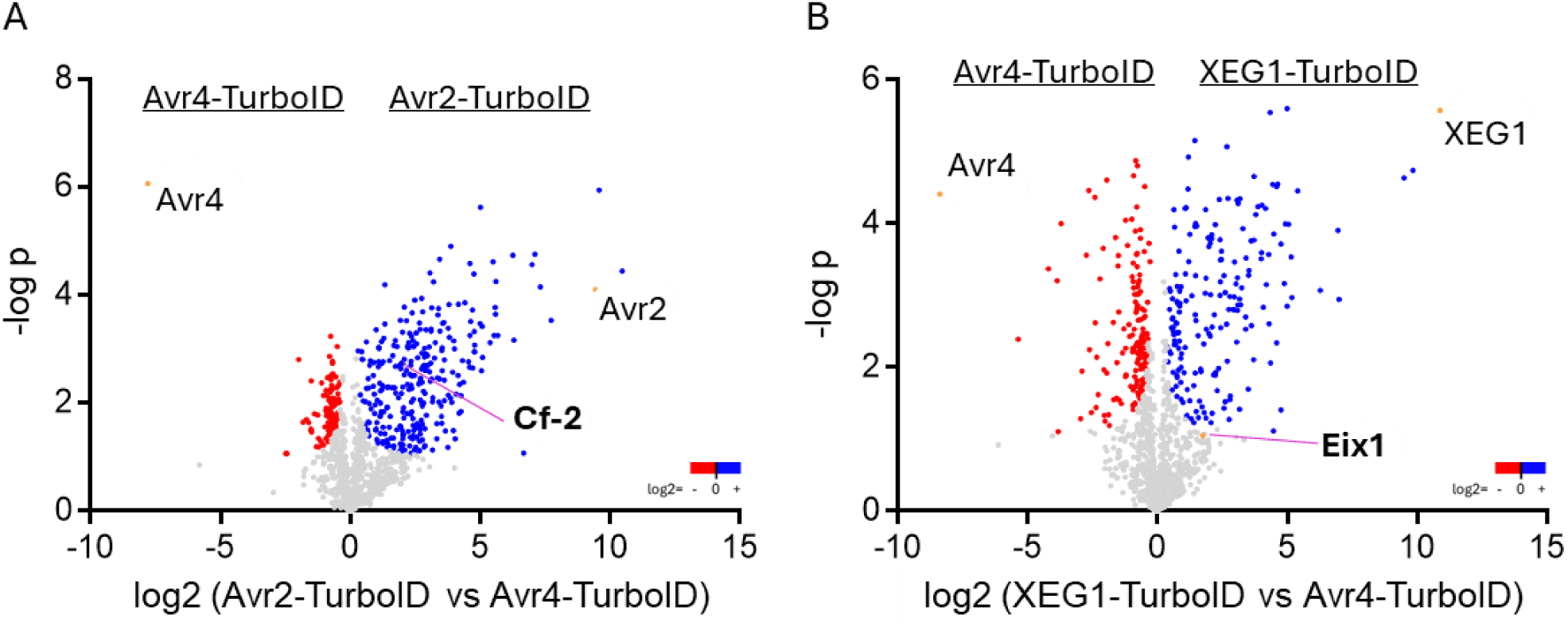
Avr2–TurboID identifies Cf-2 and XEG1–TurboID reveals *Sl*Eix1 as an XEG1-associated receptor-like protein in tomato. (A, B) Volcano plots of PL–MS data obtained following transient expression of Avr2–TurboID (A) and XEG1–TurboID (B) in leaves of MM–Cf-2 tomato plants, using expression of Avr4-TurboID as a control. Log2-transformed LFQ intensity ratios are plotted on the x-axis and −log10(*p*) values on the y-axis. Proteins significantly enriched in the Avr2–TurboID- or XEG1–TurboID-generated datasets are shown in blue while in the Avr4-TurboID-generated datasets the significantly enriched proteins are shown in red (two-sided Student’s *t*-test with permutation-based FDR < 0.05, S₀ = 0.1). Non-significantly enriched proteins are shown in grey. (A) In the Avr2–TurboID-generated samples, Cf-2 is enriched and detectable among the significantly enriched proteins, whereas Rcr3 is not detected under these conditions. (B) In the XEG1–TurboID-generated samples, the receptor-like protein *Sl*Eix1 (Q6JN47) is detected but does not pass the statistical significance threshold and therefore appears as a non-significant data point. Peptides corresponding to *Sl*Eix1 were nevertheless detected in all three biological replicates (Table S1), identifying *Sl*Eix1 as a candidate XEG1-associated RLP. All experiments were performed with three biological replicates, each prepared from five infiltrated leaves. All identified proteins upon Perseus analysis are listed in Supplementary Data 1.

### Transient expression of *Sl*Eix1 enhances the XEG1-triggered HR in a SOBIR1-dependent manner

To functionally assess the role of *Sl*Eix1 in XEG1 perception, transient co-expression assays were performed in *N. benthamiana*. For this, XEG1–TurboID was co-expressed with *Sl*Eix1 or with its close tomato homolog *Sl*Eix2 in various combinations (Figure 6). Although *N. benthamiana* harbours an endogenous functional RXEG1 receptor, a previous study has shown that transient overexpression of *Nb*RXEG1 enhances the XEG1-triggered immune response (Wang et al., 2018). By analogy, overexpression of the putative tomato ortholog *Sl*Eix1 was expected to also enhance XEG1-induced cell death. To determine whether the observed responses depend on the co-receptor SOBIR1, which has been shown to be essential for RLP functionality (Liebrand et al., 2013), the same combinations were also tested in *N. benthamiana SOBIR1* knock-out (KO) plants (Huang et al., 2021).

**Figure 6.**
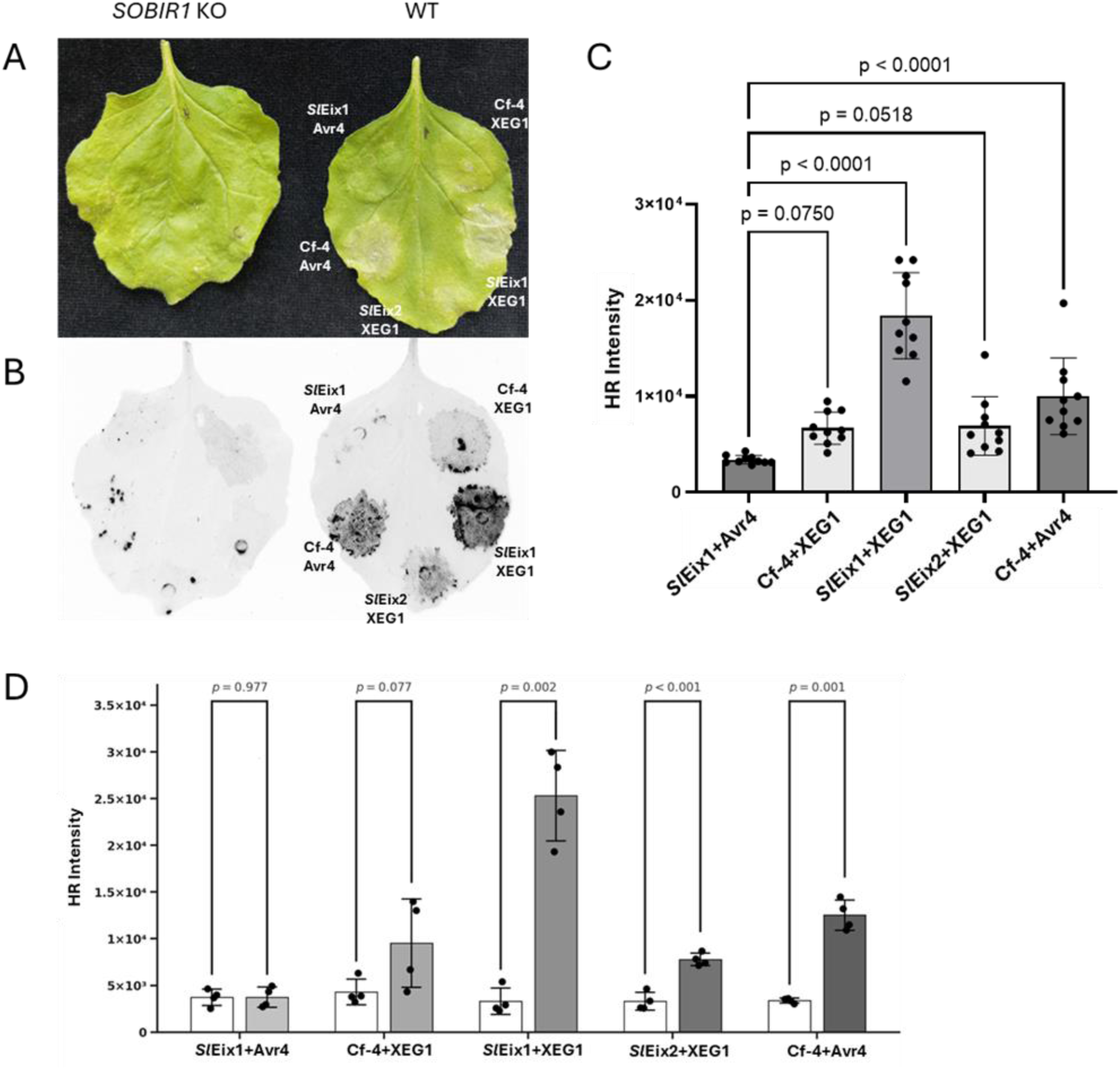
Co-expression of *Sl*Eix1 enhances the XEG1-triggered HR in a SOBIR1-dependent manner. (A, B) Representative HR phenotypes in *SOBIR1* KO and WT *N. benthamiana* leaves transiently co-expressing *Sl*Eix1, *Sl*Eix2, or Cf-4, together with XEG1–TurboID or Avr4, as indicated. Leaves were imaged at 4 dpi under visible light and by red fluorescence imaging to visualise cell death. (C) Quantification of the HR intensity in WT *N. benthamiana* based on the red fluorescence signal intensity. Bars represent mean ± SD of 10 biological replicates (infiltration spots), with individual data points shown. Co-expression of *Sl*Eix1 with XEG1–TurboID significantly increases the HR intensity. (D) Comparison of the HR intensity between the *SOBIR1* KO (white bars) and WT (grey bars) plants, triggered upon transient expression of the indicated construct combinations. Bars represent mean ± SD of four biological replicates, with individual data points shown. The HR is abolished in all cases in the *SOBIR1* KO plants, demonstrating that *Sl*Eix1-mediated enhancement of the XEG1-triggered HR requires SOBIR1. Statistical significance was assessed using one-way ANOVA followed by Tukey’s test.

In WT *N. benthamiana*, co-expression of XEG1–TurboID with *Sl*Eix1 indeed resulted in a significantly stronger and faster HR when compared with all other combinations, as assessed by visible light imaging and quantified by red fluorescence imaging (Figure 6A–C). In contrast, co-expression of XEG1–TurboID with *Sl*Eix2 did not increase the HR intensity beyond control levels, indicating the specificity of the effect of SlEix1. In the *SOBIR1* KO plants, none of the tested combinations—including *Sl*Eix1/XEG1–TurboID and the positive control Cf-4/Avr4—elicited an HR (Figure 6A, B, D). This complete loss of cell death demonstrates that the enhanced HR observed upon *Sl*Eix1 co-expression is fully dependent on SOBIR1 and therefore mediated by the RLP *Sl*EIX1.

### *Sl*Eix1 shares conserved sequence and structural features with *Nb*RXEG1 that are associated with XEG1 recognition

Given the functional association of *Sl*Eix1 with recognition of XEG1 observed in tomato and *N. benthamiana*, we next investigated whether *Sl*Eix1 shares conserved sequence motifs and structural features with the established XEG1 receptor *Nb*RXEG1. In previous studies, structural analysis of the *Nb*RXEG1–XEG1 complex has identified two receptor regions that are critical for XEG1 binding and immune activation; an N-terminal “loop-out” region (N-loopout; residues 90–95) and a C-terminal island domain (ID; residues 786–792), both of which protrude from the leucine-rich repeat (LRR) solenoid and contact the catalytic groove of XEG1 (Wang et al., 2018; Sun et al., 2022) (Figure S7). IDs are characteristic features of many LRR-containing receptor-like proteins (LRR-RLPs) and have been proposed to contribute to ligand specificity (Fritz-Laylin et al., 2005). Because *Sl*Eix1 and *Sl*Eix2 are the closest tomato homologs of *Nb*RXEG1, their amino acid sequences were aligned with *Nb*RXEG1 using Multalin (Corpet, 1988) to assess conservation of these key regions. In both the N-loopout and ID segments, *Sl*Eix1 largely retains the amino acid motifs present in *Nb*RXEG1, whereas *Sl*Eix2 displays multiple substitutions and deletions at corresponding positions (Figure S7).

To examine whether these sequence differences translate into an altered interaction potential, AlphaFold 3 (Abramson et al., 2024) models were generated for XEG1 in a complex with *Nb*RXEG1, *Sl*Eix1 or *Sl*Eix2 (Figure 7A). In the *Nb*RXEG1/XEG1 model, both the N-loopout and ID segments project towards the XEG1 surface and form part of the predicted binding interface, consistent with previously reported structural data (Sun et al., 2022). In the *Sl*Eix1/XEG1 model, these regions occupy highly similar positions, whereas in the *Sl*Eix2/XEG1 model, the ID segment appears distorted and poorly defined, which is accompanied by reduced per-residue confidence scores. While the N-loopout region remains broadly compatible with an *Nb*RXEG1-like interface in both tomato homologs, the ID segment appears structurally preserved in *Sl*Eix1 but is less conserved in *Sl*Eix2.

**Figure 7.**
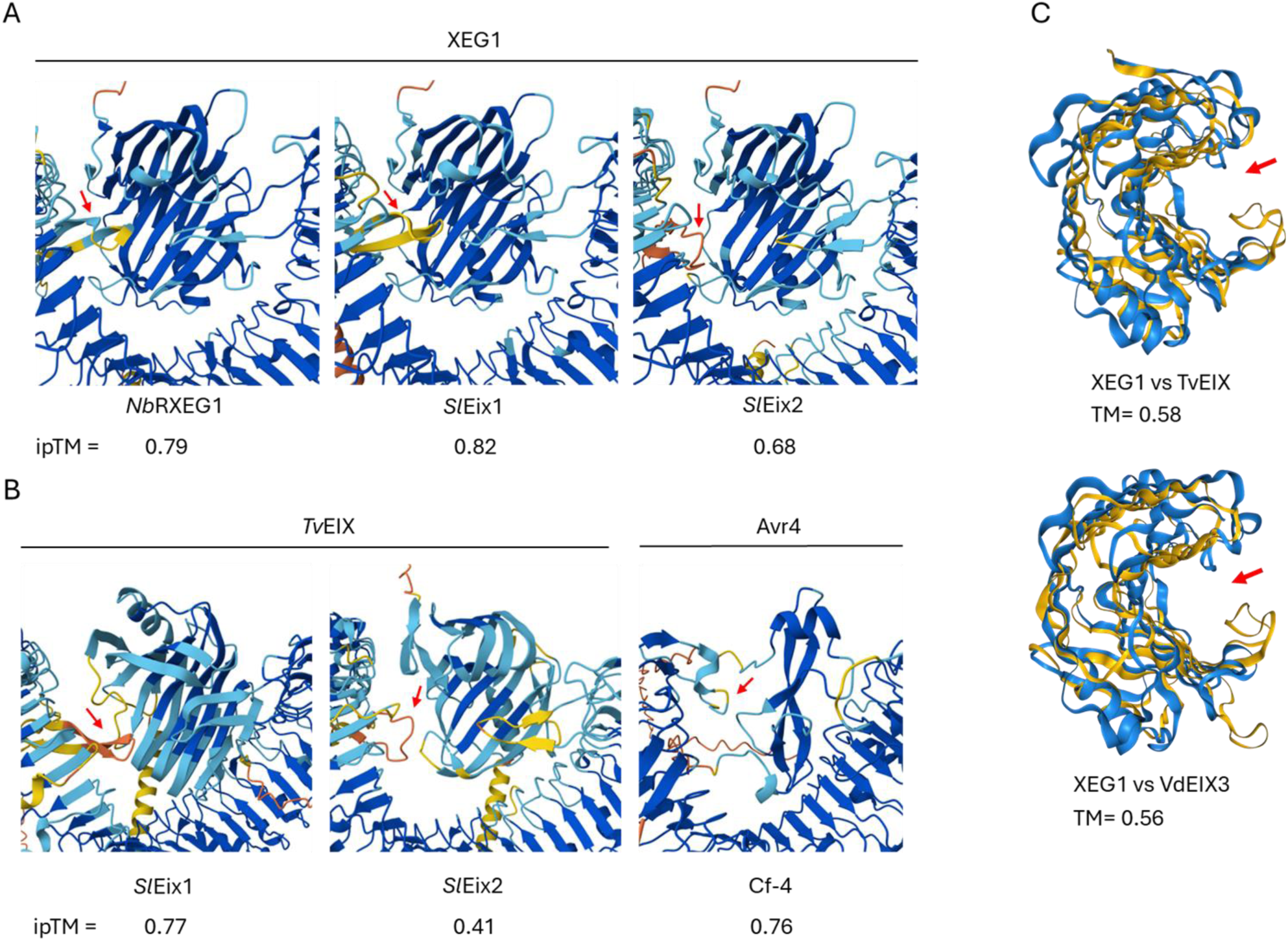
AlphaFold-based structure modelling of XEG1 and EIX interactions with *Nb*RXEG1 and tomato EIX homologs. (A) AlphaFold Multimer models of XEG1 interacting with *Nb*RXEG1 and the tomato homologs *Sl*Eix1 and *Sl*Eix2. In all models, XEG1 is positioned within the LRR solenoid. While *Nb*RXEG1 and *Sl*Eix1 display similarly positioned and well-defined ID segments, the ID of *Sl*Eix2 appears distorted and shows reduced structure confidence. Red arrows indicate the ID segments. (B) Predicted interactions between *Sl*Eix1 and *Tv*EIX, *Sl*Eix2 and *Tv*EIX, and between Cf-4 and Avr4. ipTM scores are shown for each interaction. *Sl*Eix1/*Tv*EIX and Cf-4/Avr4 interactions approach the commonly used confidence threshold of 0.8, whereas the *Sl*Eix2/*Tv*EIX interaction scores substantially lower. (C) Structural superposition of XEG1 with *Tv*EIX and *Vd*EIX3 identified by Foldseek analysis. Despite low sequence similarity, both EIX proteins exhibit significant structural similarity to XEG1 (TM scores > 0.5), including conservation of the catalytic groove (red arrow).

Consistent with these observations, the predicted interaction template modelling (ipTM) scores for *Sl*Eix1/XEG1 and *Nb*RXEG1/XEG1 interactions are close to the commonly used confidence threshold of 0.8, whereas the *Sl*Eix2/XEG1 interaction scored an ipTM that is substantially lower (0.68) (Figure 7B). Interestingly, modelling of *Sl*Eix1 interacting with the *Trichoderma viride* elicitor *Tv*EIX yielded an ipTM score comparable to those of the *Sl*Eix1/XEG1 (0.77) and Cf-4/Avr4 (0.76) interactions, while the *Sl*Eix2/*Tv*EIX interaction resulted in a markedly lower score (0.41). Given the apparent ability of *Sl*Eix1 to associate with both XEG1 and EIX-type elicitors, we examined whether these elicitors share structural similarity. A Foldseek-based structural similarity search (van Kempen et al., 2024), using the XEG1 3D structure (PDB ID: 7DRC), identified *Tv*EIX and *Verticillium dahliae Vd*EIX3 as structurally related proteins, with template modelling (TM) scores of 0.58 and 0.56, respectively, exceeding the commonly accepted threshold of 0.5 for structural similarity (Xu et al., 2010) (Figure 7C). This similarity was observed despite having a low primary sequence identity (Figure S8), suggesting that conserved three-dimensional features, including the catalytic groove, may underlie the ability of *Sl*Eix1 to associate with both XEG1 and EIX-type elicitors.

## DISCUSSION

Proximity-dependent labelling (PL) has become a very powerful strategy to map protein interaction networks *in vivo* and to dissect the composition of immune receptor complexes across eukaryotes (Roux et al., 2012; Branon et al., 2018; Zhang et al., 2019; Huang et al., 2024). In plants, TurboID-based PL has so far been used mainly to identify host factors associated with pathogen effectors, disease susceptibility processes, and downstream signalling modules, rather than to systematically connect apoplastic effectors to their corresponding cell-surface R proteins. Recent examples include effector-centred PL in maize (*Zea mays*), where TurboID fused to a *Puccinia polysora* effector identified MITOGEN-ACTIVATED PROTEIN KINASE 3 (MPK3) as a susceptibility factor (Su et al., 2025), and PL approaches that defined interaction partners of viral effectors in *N. benthamiana* and various crop plants (Cao et al., 2010; Navarro et al., 2021; Sáiz-Bonilla et al., 2025). Effector–TurboID fusions have also been applied to identify targets of nematode and bacterial effectors, including the association between the nematode effector *MELOIDOGYNE INCOGNITA*-SPECIFIC PIONEER 18 (MSP18) and the receptor-like cytoplasmic kinase BRASSINOSTEROID SIGNALING KINASE 7 (BSK7) and between the RALSTONIA INJECTED PROTEIN U (RipU) and the RPM1-INTERACTING PROTEIN 4 (RIN4) (Chen et al., 2025; de Ryck et al., 2025). In parallel, TurboID fused to NLRs has been used to define immune signalling modules and paired NLR behaviour *in planta* (Zhang et al., 2019; Zhang et al., 2024; Chen et al., 2025), and a complementary “reverse” strategy has been proposed in which NLR-linked TurboID supports the discovery of matching effectors and associated host components (Szymansky, 2025). Collectively, these studies demonstrate the versatility of PL in plant immunity research, but the studies also highlight a remaining gap: being practical methods that are able to match apoplastic effectors to PM-localised PRRs, which can be either RLPs or RLKs, in crop systems.

Here, we establish effector–TurboID-mediated PL as a practical route to identify PM-associated receptors that recognise apoplastic effectors in *N. benthamiana* and tomato. By fusing three apoplastic effectors (Avr2, Avr4 and XEG1) to TurboID and transiently expressing the protein fusions in leaves, we show that apoplastic PL can recover receptor proximity signatures representing both direct and indirect recognition systems *in planta*. We have validated the approach using the Cf-4/Avr4 and Cf-2/Avr2 pairs (Dixon et al., 1996; Thomas et al., 1997), and have applied it to XEG1, for which the corresponding tomato receptor was unknown at the start of this work (Wang et al., 2018).

### Functional validation of effector–TurboID fusions

A prerequisite for apoplastic PL is that large effector–TurboID fusion proteins traverse the secretory pathway, are properly folded, accumulate extracellularly, and retain sufficient biological activity. Immunoblotting of apoplastic fluids from agroinfiltrated *N. benthamiana* leaves confirmed the accumulation of full-length Avr2–, Avr4–, and XEG1–YFP–TurboID fusions in the apoplast (Figure S3). Avr2–YFP–TurboID accumulated to relatively high levels in the AF, which is consistent with Avr2 acting as a protease inhibitor that stabilises itself and potentially other apoplastic proteins (Rooney et al., 2005; van Esse et al., 2008; Shabab et al., 2008). In line with this, co-expression of the apoplastic protease inhibitor EPI1 increased the accumulation of Avr4–YFP–TurboID, supporting the view that extracellular proteolysis contributes to at least partial processing of these fusions (Figure S1).

Catalytic competence at the cell surface was supported by streptavidin enrichment assays: Cf-4–Myc and *Nb*RXEG1–Myc were recovered in a co-IP only when the corresponding effector–TurboID fusion was present (Figure 1C and Figure S2), whereas HR assays provided an additional functional readout. Indeed, Avr4-TurboID triggered a Cf-4-dependent HR comparable to its untagged counterpart, indicating that the TurboID fusion does not measurably compromise the presumed direct recognition of Avr4 by Cf-4. In contrast, Avr2-TurboID induced a weaker HR than the non-fused protein in combination with Cf-2, which, considering the indirect recognition, suggests that the fusion modestly reduces the signalling efficiency, without abolishing recognition (Figure S4). Overall, these tests establish that effector–TurboID fusions are secreted, remain catalytically active, and retain sufficient biological activity to serve as PL baits.

Furthermore, Avr4 was expressed using its native fungal signal peptide for extracellular targeting, whereas Avr2 and XEG1 employed the *Nb*PR1a signal peptide, which is widely used for efficient extracellular targeting in Solanaceae (Agarwal et al., 2008; Kanagarajan et al., 2008; Mesarich et al., 2014). KEGG enrichment analyses of the effector–TurboID proxitomes reveal that all three fusions cause abundant enrichment of translation-related categories, as expected for highly expressed proteins, whereas apoplast- and secretory pathway-related terms are exceptionally prominent for Avr2 and XEG1, which make use of the NbPR1a signal peptide. (Figure S9).

Finally, extracellular ATP infiltration consistently increased overall protein biotinylation levels in tomato (Figure 3). In addition to its role in enzymatic reactions like biotinylation, extracellular ATP is a well-established DAMP that is recognised by the legume-type lectin receptor kinase DOES NOT RESPOND TO NUCLEOTIDES 1 (DORN1) and related receptors, and thereby ATP boosts immune signalling and might increase the abundance or activation state of immune-related RLKs and/or RLPs at the PM (Choi et al., 2014; Saijo & Loo, 2018; Vega-Muñoz et al., 2020). Notably, ATP addition coincided with stronger Cf-4 enrichment in tomato when compared to *N. benthamiana*, despite higher Avr4–TurboID accumulation in the latter plant, suggesting that ATP supplementation may indeed improve receptor detection by both enhancing catalysis by the TurboID enzyme and favouring immune complex availability.

### Endogenous receptor identification in *N. benthamiana* and tomato

Following validation, effector–TurboID PL–MS recovered endogenous receptors in both model and crop backgrounds. In *N. benthamiana:Cf-4*, Avr4– and XEG1–TurboID proxitomes were enriched for Cf-4 and NbRXEG1, respectively (Figure 2), demonstrating that effector-centred PL is able to identify RLP-based recognition at the protein level. Importantly, these receptors were detected without their constitutive overexpression, as Cf-4 is expressed under its native promoter in the transgenic *N. benthamiana* line, and *NbRXEG1* is an endogenous gene, indicating that PL captures receptor proximity signatures even at relatively modest expression levels.

Transfer of the approach to the tomato near-isogenic introgression lines provided a stringent test in a crop host. In MM–Cf-4 plants, Avr4–TurboID robustly enriched Cf-4 in PL–MS datasets, consistent with direct ligand binding to the Cf-4 receptor at the cell surface (Figure 4). In MM–Cf-2 plants, Avr2–TurboID enriched Cf-2 among significantly enriched proteins, although Cf-2 was not an extreme volcano-plot outlier (Figure 5). Interestingly, Rcr3 was not enriched in this background, whereas Avr2–TurboID did enrich Rcr3 and additional apoplastic proteases in MM–Cf-4 plants (Figure 4). This reciprocal pattern is consistent with known relationships between Avr2, its virulence target Rcr3, and receptor-mediated perception (Kourelis et al., 2024), and suggests that an effector–TurboID fusion recovers both guarded effector targets and receptors depending on the receptor context.

XEG1 presented the most challenging case because the corresponding tomato receptor was not established initially. In MM–Cf-2 tomato, XEG1–TurboID did not yield an RLP that exceeded the standard significance threshold. However, closer inspection revealed one RLP (Q6JN47) that was detected exclusively in the XEG1–TurboID-generated samples across all three replicates but was not classified as significant due to low peptide counts in one replicate. Q6JN47 corresponds to *Sl*Eix1 (Solyc07g008620), which has previously been proposed to act primarily as a decoy that attenuates EIX responses (Bar et al., 2010). The replicate-consistent presence of *Sl*Eix1 peptides in the XEG1–TurboID-generated datasets (Table S1) supports *Sl*Eix1 as a biologically meaningful, low-abundance candidate receptor. Functional tests supported this interpretation. In *N. benthamiana*, co-expression of XEG1–TurboID with *Sl*Eix1, but not *Sl*Eix2, indeed enhanced the XEG1-triggered HR in a SOBIR1-dependent manner. Together with the PL–MS data, these data are consistent with *Sl*Eix1 functioning as the tomato ortholog of *Nb*RXEG1 in XEG1 perception. Recently, independent support for our observation was provided by Zeng et al. (2025), who reported that *Sl*Eix1 complements *Nb*RXEG1 function and restores the XEG1-triggered HR in an *rxeg1* mutant of *N. benthamiana*. Sequence and structure comparisons further align with this model. *Sl*Eix1 is more similar to *Nb*RXEG1 than *Sl*Eix2 in regions implicated in XEG1 binding and receptor activation, including the N-loopout and ID segments (Wang et al., 2018; Sun et al., 2022) (Figure 7 and Figure S7). While the N-loopout is broadly conserved in both paralogs, *Sl*Eix1 retains an *Nb*RXEG1-like ID with higher-confidence interaction features, whereas *Sl*Eix2 is divergent and associated with reduced AlphaFold-derived interaction confidence. These data are consistent with a model in which *Sl*Eix1, but not *Sl*Eix2, preserves an *Nb*RXEG1-like recognition interface for XEG1, while *Sl*Eix1 may also retain the capacity to associate with EIX-like elicitors.

### The tomato *EIX* locus as a multi-effector recognition hub

The tomato *EIX* locus was originally defined by recognition of the *T. viride* elicitor EIX (Ron & Avni, 2004). Two RLP-encoding genes, *SlEix1* and *SlEix2*, were first cloned from this locus; *Sl*Eix2 mediates the EIX-triggered HR, whereas *Sl*Eix1 was found to bind EIX without triggering an HR and, when overexpressed, attenuates *Sl*Eix2-mediated responses (Bar et al., 2010). Beyond *Tv*EIX, *Vd*EIX3 also activates Eix2-dependent responses (Yin et al., 2021). The *EIX* locus contains additional genes encoding EIX-LIKE (Eil) proteins; notably, Solyc07g008600 (*Sl*Eil2/*Sl*8600) functions as the receptor for the *Sclerotinia sclerotiorum* SMALL CYSTEINE-RICH SECRETORY PROTEIN (SsSCP) (Zhang et al., 2025). Our PL and functional data, together with recent evidence that *Sl*Eix1 recognises multiple GH12 proteins, including XEG1 (Zeng et al., 2025), extend this locus-level view by positioning *Sl*Eix1 as an XEG1-associated receptor component.

The *EIX* locus therefore encodes receptors for at least three distinct effectors from different fungal and oomycete pathogens, being EIXs (GH11 family glycoside hydrolases), *Tv*EIX and *Vd*EIX3, *Ss*SCP (an unclassified small secreted cysteine-rich protein) and 15 GH12 family glycoside hydrolases from various pathogens, including XEG1 itself (Figure 8). This functional diversification within a single RLP-encoding cluster resembles other complex *R* gene loci, such as the *homologs of Cladosporium-resistance gene Cf-9 at the Cf-9 locus* (*Hcr9-9s*) and those at the *Cf-4* locus (*Hcr9-4s*), containing *Cf-9*/*Cf-9B* and *Cf- 4*/*Cf-4E*, respectively, encoding receptors with distinct effector specificities (Thomas et al., 1997; Takken et al., 1999; de la Rosa et al., 2024). However, the *EIX* locus is unusual in its breadth (Figure S10) and taxonomic diversity of the recognised ligands (Figure S11).

**Figure 8.**
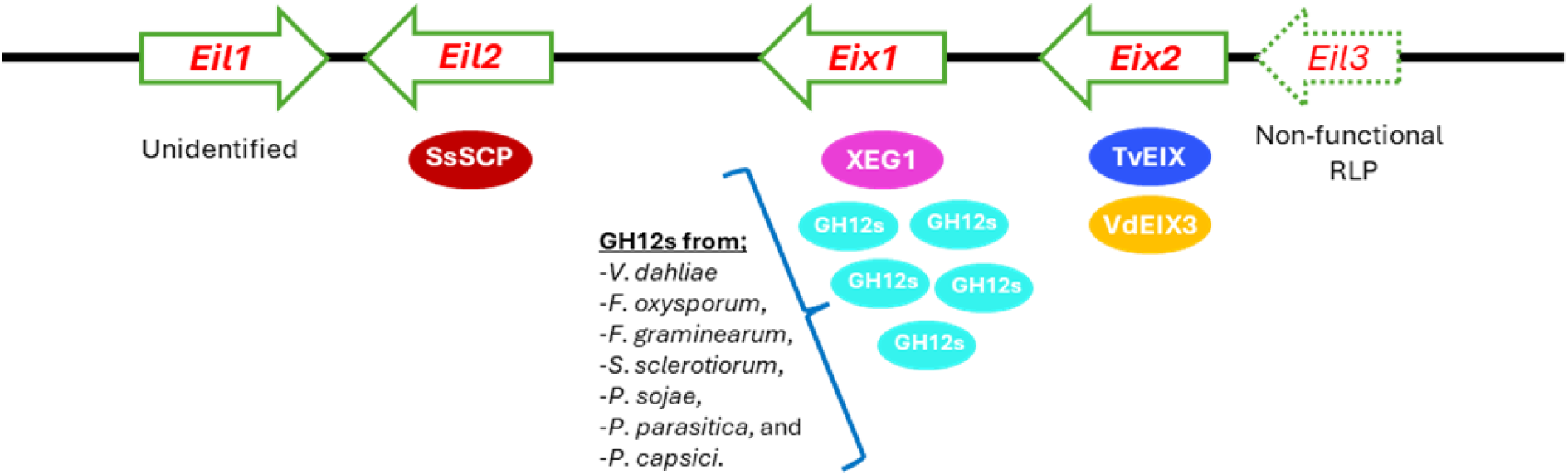
Genomic organisation of the tomato *EIX* locus and an overview of its matching effectors. Schematic representation of the *EIX* locus located on chromosome 7, showing the relative positions and transcriptional orientation of *Eil1* to *Eil3*, *SlEix1* and *SlEix2*. Below the scheme, known and proposed effector specificities of the encoded receptors are indicated; *Eil2* encodes the primary receptor for SsSCP, *Eix2* encodes the primary receptor for *Tv*EIX/*Vd*EIX3, and *Eix1* encoding the receptor for XEG1 and other GH12 proteins. This locus illustrates the organization of a multi-effector immune hub, comprising both signalling-competent and possibly decoy-type RLPs.

### *Sl*Eix1 versus *Sl*Eix2: context-dependent signalling and regulation

Recent work suggests that *Sl*Eix1 also acts as a negative regulator, as knocking out *SlEix1* enhances resistance and is associated with altered expression of other PRRs (Leibman-Markus et al., 2023), potentially reflecting resource allocation or compensatory rewiring in receptor biogenesis and immune signalling. In apparent contrast, Zeng et al. (2025) conclude that *Nb*RXEG1 and its Solanaceae orthologs, including *Sl*Eix1, contribute to broad-spectrum resistance by recognising multiple microbial GH12 proteins. In *N. benthamiana*, combined loss of both *NbEix2* and *NbRXEG1* abolishes the EIX-triggered HR, whereas single mutants do not, suggesting that *Nb*RXEG1 contributes to EIX perception in addition to its ability to recognize XEG1 (Wang et al., 2018). Together with the observation that physical interaction takes place between *Sl*Eix1 and *Sl*Eix2 (Bar et al., 2010), these findings support a model in which heterocomplex formation among the EIX-locus-encoded RLPs may contribute to perception of EIX-like elicitors, potentially with functional redundancy among the locus members in tomato. The observation that *Sl*Eix1 overexpression attenuates the EIX-mediated HR (Bar et al., 2010), underscores context and dosage sensitivity. A similar dosage-sensitive behaviour has been reported for the helper NLR of tomato NB-LRR REQUIRED FOR HR-ASSOCIATED CELL DEATH 4a (*Sl*NRC4a), which interacts with *Sl*Eix2 and triggers an increased ROS production upon EIX perception when either overexpressed or knocked out (Leibman-Markus et al., 2018).

Cf-4 and *Sl*Eix2 have overall structural similarity, and for both of them at the C-terminus a conserved Yxxϕ endocytosis motif is present (Sharfman et al., 2011; Postma et al., 2016), identical to Cf-9 (Thomas et al., 1998), and they both undergo ligand-triggered endocytosis (Bar & Avni, 2010; Chakrabarti et al., 2016; Postma et al., 2016). However, helper NLR dependencies differ, as Cf-4 responses depend on *Sl*NRC1/2/3 but not on *Sl*NRC4 (Gabriëls et al., 2007; Kourelis et al., 2022), whereas *Sl*NRC4a impacts *Sl*Eix2-mediated outputs as mentioned above (Leibman-Markus et al., 2018). Together, available data support a scenario in which EIX perception involves receptor complexes with distinct co-receptor/helper wiring and quantitative constraints, rather than a strictly linear pathway. Resolving the composition and dynamics of these complexes remains an important goal of our research on plant immunity.

### Effector–TurboID-based PL as part of a broader *R* gene discovery pipeline

Effector–TurboID PL complements sequence-based cloning methods such as RenSeq, MutRenSeq and AgRenSeq, which have accelerated *R* gene discovery, but largely operate at the DNA level (Jupe et al., 2013; Steuernagel et al., 2016; Arora et al., 2019; Zhang et al., 2020; Lin et al., 2020). This complementarity is particularly valuable for crop species in which genetics-based *R* gene discovery remains challenging, including polyploid genomes with extensive redundancy and structural variation, as well as perennial crops with long generation times that preclude rapid population development and fine-mapping. By interrogating the protein complexes assembled around an effector directly *in planta*, effector-centred PL–MS bypasses these constraints and captures not only primary receptors but also co-receptors, adaptors and regulatory proteins, including decoys and guarded host targets.

Methodologically, our results also suggest introducing practical improvements in future screens. DAMP priming, for example, by the addition of extracellular ATP, and optimised signal peptide preference will enhance receptor labelling. In addition, using stronger extraction buffers, such as radio-immunoprecipitation assay RIPA buffer (Ngoka, 2008), may increase the recovery of membrane-associated proteins such as RLPs (Liebrand et al., 2012) and replacing YFP with a smaller, flexible linker may reduce steric interference and be especially beneficial for dissecting indirect recognition systems such as Cf-2/Rcr3/Avr2. Moreover, integration of the data analysis with long-read full-length transcript sequencing platforms (Iso-Seq) will facilitate accurate reconstruction of complete receptor-coding sequences and splice variants, particularly in crops with complex or poorly annotated genomes, thereby accelerating downstream cloning and functional validation.

In summary, effector–TurboID-mediated PL enables (i) recovery of known RLP-based resistance systems, (ii) identification of new functional roles for previously characterised receptors within complex loci, and (iii) mechanistic dissection of how closely related receptors partition effector recognition. By directly linking pathogen effectors to their corresponding cell-surface receptors at the protein level, this approach substantially shortens the path from pathogen genome to a deployable *R* gene. Although demonstrated here for extracellular effectors and cell-surface immune receptors, the same conceptual framework should also be applicable to cytoplasmic effectors and intracellular immune receptors. Effector-centred PL therefore provides a powerful complement to existing genetic and sequence-based strategies, with particular promise for accelerating resistance breeding in crops with complex genomes, limited genetic resources, and/or long breeding cycles. More broadly, this PL approach opens new opportunities to discover and exploit immune receptors in previously underexplored or recalcitrant crop species, thereby expanding the repertoire of *R* genes available for durable disease control.

## MATERIALS AND METHODS

### Plant material and growth conditions

*N. benthamiana* WT, *N. benthamiana SOBIR1* KO, and *N. benthamiana:Cf-4* plants were grown in climate chambers, and tomato Moneymaker (MM)-Cf-4 and MM-Cf-2 lines were grown in a greenhouse, all under 16 hr of light at 24 °C, and 8 hr of darkness at 22 °C, and at a relative humidity of 75%.

### Generation of binary vectors and Agrobacterium-mediated transient transformation

The constructs used for the PL-MS experiments, Avr2-YFP-TurboID, Avr4-YFP-TurboID and XEG1-YFP-TurboID were made in the following way. The coding sequence corresponding to the 135 amino acid primary translation product of the *Avr4* gene was amplified from the pMOG800-Avr4 (SOL6783) plasmid. The *Avr2* coding sequence, in which the native signal peptide for extracellular targeting was replaced by the signal peptide encoded by the *NbPR1a* gene, was amplified from the pSfinx::PR1a-Avr2 (SOL10085) plasmid. Both inserts were then inserted separately into the linearised pEG101-YFP–TurboID (SOL6890) vector. For the construction of the XEG1 fusion, the coding sequence of *XEG1*, excluding the first 54 nucleotides corresponding to its native signal peptide, was amplified from *P. sojae* cDNA. The *NbPR1a* signal peptide was amplified from the pSfinx::PR1a-Avr2 plasmid, and the *PR1a* and *XEG1* fragments were subsequently cloned into the linearised pEG101-YFP–TurboID vector. All DNA fragments were amplified using Phusion Hot Start II DNA Polymerase (Thermo Scientific) and assembled using the ClonExpress MultiS One Step Cloning Kit (Vazyme). The nucleotide sequences of the primers used for all DNA amplifications are listed in Table S2.

Agroinfiltration was performed as previously described (van der Hoorn et al., 2000). We performed agroinfiltration in the first fully expanded leaves of three- to four-week-old *N. benthamiana* and tomato plants at an OD_600_ of 0.5 and 0.1, respectively. The lower OD was used in tomato with the aim to minimise infiltration-induced necrosis. Each sample derived from leaves expressing one of the TurboID-tagged fusion proteins was prepared in three biological replicates. For each replicate, we infiltrated one leaf per plant, with a total of five plants. The leaves for generating the samples were randomly assigned to different plants.

### Apoplastic fluid (AF) isolation

Apoplastic fluid (AF) was isolated from infiltrated *N. benthamiana* leaves at 1–2 days after agroinfiltration as described previously (Joosten, 2012), with minor modifications. Five infiltrated leaves were submerged in deionised water in a 500 mL beaker and gently weighted down to prevent flotation. The beaker was placed in a desiccator and leaves were vacuum-infiltrated at ≤60 mbar for 5 min to facilitate infiltration of the apoplastic space. Following the infiltration, leaves were briefly blotted dry, rolled and transferred to a 20 mL syringe without a plunger, which was positioned inside a 50 mL centrifuge tube. AF was recovered by centrifugation at 3,000 × g for 5 minutes (min) at 4°C. AF samples were directly mixed with SDS loading buffer (200 mM Tris-HCl, pH 6.8, 8% SDS, 40% glycerol, 400 mM DTT, 2% bromophenol blue), boiled at 95 °C for 5 min, and subjected to SDS–PAGE and immunoblot analysis as described below.

### Total protein extraction and streptavidin affinity purification

At 2-3 days after agroinfiltration, leaves were infiltrated with a solution containing ATP (1 mM + 5 mM MgCl_2,_ which is required for proper functioning of ATP (Campos & Beaugé, 1992)) and biotin (200 μM, pH 8, in 10 mM MES buffer) and harvested 1 hour later. For each replicate, the five treated leaves were harvested, combined, wrapped in aluminium foil, and kept in liquid nitrogen or at −80 °C until further use. The frozen samples were ground to a fine powder in liquid nitrogen using a mortar and pestle. Then, the pulverised samples were transferred to pre-cooled 50 mL V-shaped tubes, weighed, and resuspended in 2 mL/g of extraction buffer (EB; pH = 8.0, 150 mM NaCl, 1.0% [v/v] IGEPAL^®^ CA-630 (NP-40), 50 mM Tris, and Sigma protease inhibitor cocktail (1 tablet per 50 mL)). For resuspension, the frozen samples with EB were vortexed at room temperature and kept on ice after melting. Then, the samples were centrifuged at 19,650 x g for 45 min at 4°C in a Sigma 4-16K centrifuge.

To prevent saturation of the streptavidin beads with the free biotin remaining in the samples, 10 mL of the centrifuged total protein extracts were desalted using PD MiniTrap PD-10 desalting columns (GE Healthcare) at 4 °C, following the manufacturer’s gravity protocol for salt removal. The desalted protein extracts were then transferred to pre-cooled 15 mL tubes and incubated for 1 hour at 4 °C (10 rpm on an SB3 tube rotator, STUART) with 200 μL of Dynabeads™ MyOne™ Streptavidin C1 (washed before use according to the manufacturer’s protocol). After incubation, the beads were washed three times with 1 mL of EB (without NP-40) and resuspended in 45 μL of EB. After the final EB wash, samples were either processed for immunoblot analysis or subjected to on-bead tryptic digestion for LC–MS/MS.

### Immunoprecipitation (IP)

The accumulation, receptor interaction and biotinylation activity of the TurboID fusion proteins were examined by protein IP or co-IP analysis, followed by immunoblotting as described previously (Liebrand et al., 2012). After the final wash step, beads were resuspended in SDS loading buffer and boiled at 95 °C for 5 min. Eluted proteins were separated by SDS–PAGE and transferred to PVDF membranes using a Trans-Blot Turbo system (Bio-Rad) (1.3 A, 25 V, 7 min). Membranes were blocked in TBS-T (150 mM NaCl, 20 mM Tris-HCl, pH 7.5, 0.1% Tween-20), containing 5% (w/v) skimmed milk for 1 h at room temperature or overnight at 4 °C, unless stated otherwise.

For detection of TurboID fusion proteins, membranes were incubated with rabbit anti-BirA (mutated TurboID) antibody (Agrisera) at a 1:5,000 dilution, followed by goat anti-rabbit HRP-conjugated secondary antibody (Bio-Rad; 1:10,000). Biotinylated proteins were detected using a streptavidin–HRP conjugate (GE Healthcare; 1:2,000). For streptavidin–HRP detection, membranes were blocked in TBS-T containing 5% (w/v) BSA instead of using milk.

### Sample Preparation for Proteomics by LC-MS/MS

After the final washing step with EB, the beads were subjected to three additional washing steps with 1 ml of ABC buffer (50 mM ammonium bicarbonate, pH=8) and were eventually resuspended in 45 μL of ABC buffer. While still on the beads, the disulfide bonds in the captured proteins were reduced by adding 5 μL of DTT (150 mM) and incubating the samples at 45°C for 30 min. The sulfhydryl groups were subsequently alkylated by adding 6 μL of acrylamide (200 mM) and incubating the samples for 10 min at room temperature.

The peptides to be measured by LC-MS/MS were subsequently released from the streptavidin-coated beads by tryptic digestion. For this, a stock solution of trypsin (0.5 μg/μL of trypsin in 1 mM HCl, pH 3) was diluted 100 times in ABC buffer, of which 100 μL was added to each sample. The samples were then incubated overnight at room temperature, with mild agitation, after which they were acidified to pH=3 using trifluoroacetic acid, and cleaned up using μColumns according to the method described previously (Wendrich et al., 2017).

### LC-MS/MS Analysis

For the LC-MS/MS analysis, the peptides were separated by reverse-phase nano liquid chromatography using a Thermo nLC1000 column, after which they were measured using an Orbitrap Exploris 480 mass spectrometer. The peptide spectra were searched in MaxQuant (version 2.0.3.0) (Cox & Mann, 2008), using the Andromeda search engine (Cox et al., 2011) with label-free quantification (LFQ), against the Niben1.0.1 version of the *N. benthamiana* or tomato proteome datasets, including the protein sequence of Cf-4 (O50025), Cf-2 (Q41398), XEG1 (G4ZHR2), Avr4 (Q00363), Avr2 (Q8NID8) and frequently occurring contaminants.

The identified protein groups were then analysed using Perseus (version 1.6.2.3) (Tyanova et al., 2016). Reverse and contaminant proteins, as well as those identified only by matching, were filtered out. Also, protein groups identified in fewer than three replicate samples were filtered out. The LFQ values were log2-transformed, and the missing values were assigned assuming a normal distribution. Protein quantitation of the samples relative to each other was calculated by applying two-sided Student’s t tests, using a permutation-based adjustment (FDR=0.05, 250 randomisations, and S0 set to 0.1).

### AlphaFold-based structural modelling and Foldseek analysis

The 3D structure models used in this study were generated via the Alphafold Server (Abramson et al., 2024). Full-length sequences of the protein were used for the analysis. Top-ranked models were selected. The 3D structure of XEG1 (PDB ID: 7DRC) was used as a search template via the Foldseek Search Server. The search was made on the PDB100 database. Structural models were used for hypothesis generation and interpretation of experimental data, not as standalone evidence of interaction.

## AUTHOR CONTRIBUTIONS

NC, EB, SLV, CRS, and MHAJ designed the research; NC, EB, SLV, CRS, CTW, KE, and SB performed the research; NC, EB, SLV, and SB carried out the data analysis; NC, CRS, and MHAJ wrote the manuscript. All authors reviewed the manuscript and approved it for publication.

## ACKNOWLEDGMENTS

We acknowledge the personnel of UNIFARM, especially Bert Essenstam, for excellent plant care and Renier van der Hoorn for providing the binary *EPI1* construct. NC and EB are supported by the Ministry of National Education of Türkiye. SLV is supported by the Peruvian Council for Science, Technology and Technological Innovation (CONCYTEC) and its executive unit, FONDECYT.

## CONFLICTS OF INTEREST

NC, CRS, SLV, and MHAJ are inventors of the patent application: Methods of identifying plant resistance proteins, under the patent number WO2025095780.

## DATA AVAILABILITY STATEMENT

The data that support the findings of this study are available in the Supplementary Material of this article.

## SUPPLEMENTARY MATERIAL (brief legends)

**Figure S1.** Co-expression of the protease inhibitor EPI1 increases the accumulation of the Avr4-YFP-TurboID fusion protein in the apoplast.

**Figure S2.** XEG1–TurboID biotinylates its matching receptor RXEG1 upon their transient co-expression in leaves of *N. benthamiana*.

**Figure S3**. Apoplastic accumulation of effector–YFP–TurboID fusions upon their transient expression in leaves of *N. benthamiana*.

**Figure S4.** Comparison of the intensity of the HR induced by Avr2 and Avr4, with or without a TurboID fusion.

**Figure S5.** Principal component analysis of effector–TurboID PL–LC-MS/MS datasets.

**Figure S6.** Effector–YFP–TurboID fusion proteins are properly expressed in tomato leaves.

**Figure S7.** Amino acid sequence alignment of *Nb*RXEG1 and its closest tomato orthologs, *Sl*Eix1 and *Sl*Eix2.

**Figure S8.** Amino acid sequence alignment of XEG1, *Tv*EIX and *Vd*EIX3.

**Figure S9.** GO and pathway enrichment of the effector–TurboID proxitomes in (A) *N. benthamiana* and (B) tomato.

**Figure S10.** Alignment of the amino acid sequences of the RLPs encoded by the tomato *EIX* locus.

**Figure S11.** Phylogenetic relationships between the RLPs encoded by the *EIX* locus and the Cf-2 and Cf-4 proteins.

**Figure S12.** Peptide coverage of Cf-4, Cf-2, and Eix1 in the PL–LC-MS/MS datasets generated from tomato.

**Table S1.** Number of *Sl*Eix1-derived peptides detected in each replicate of the LC-MS/MS data generated by transiently expressing the different effector-TurboID fusions in leaves of MM-Cf-2 tomato plants.

**Table S2.** Nucleotide sequences of the primers used in this study.

**Supplementary Data 1.** Proteins identified in effector-TurboID samples following MaxQuant and Perseus analysis. UniProt identifiers were used for ID mapping to assign Solyc gene IDs, Gene Ontology terms, and protein family annotations in tomato. *N. benthamiana* gene annotations were directly added using Perseus.

