## Supplementary material for "Effector-centred proximity-dependent labelling enables the discovery of cell-surface immune receptors in plants": Figures S1-S12, Tables S1-S2

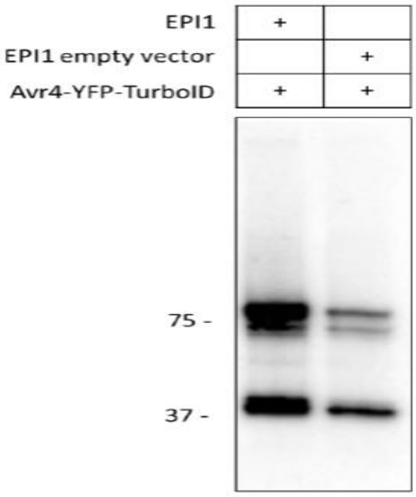


**Figure S1.** Co-expression of the protease inhibitor EPI1 increases the accumulation of the Avr4-YFP-TurboID fusion protein in the apoplast. *N. benthamiana* plants were either co-coagroinfiltrated with EPI1 or the empty vector of EPI1, and with Avr4-YFP-TurboID. At 2 dpi, apoplastic fluid was isolated and ~11 µl was run on an SDS-PAGE gel. After WB, αTurboID antibodies were used to visualise the Avr4-YFP-TurboID fusion protein. Note that co-expression with EPI1 results in overall higher accumulation levels of the recombinant Avr4-YFP-TbID fusion protein at 80kDa, and also of its degradation products.

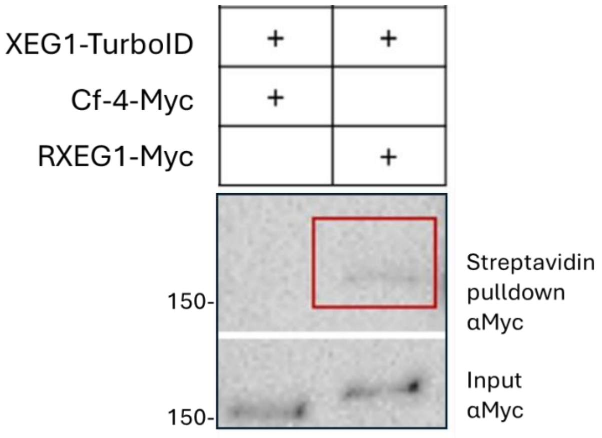


**Figure S2.** XEG1–TurboID biotinylates its matching receptor RXEG1 upon their transient co-expression in leaves of *N. benthamiana*. Leaves of WT *N. benthamiana* plants were co-infiltrated with XEG1–YFP–TurboID and *Nb*RXEG1-Myc or with non-matching Cf-4-Myc, as indicated. At 2 dpi, a 200μM biotin solution was infiltrated, and leaves were incubated for 1 h before protein extraction. Total protein extracts were subjected to streptavidin pulldown and analysed by WB using anti-Myc antibodies. *Nb*RXEG1-Myc, but not Cf-4-Myc, was biotinylated by XEG1–YFP–TurboID, as *Nb*RXEG1-Myc is captured by the streptavidin pulldown. The input (total protein extract) confirms expression of both Cf-4-Myc and NbRXEG1-Myc. Molecular mass markers (kDa) are indicated on the left.

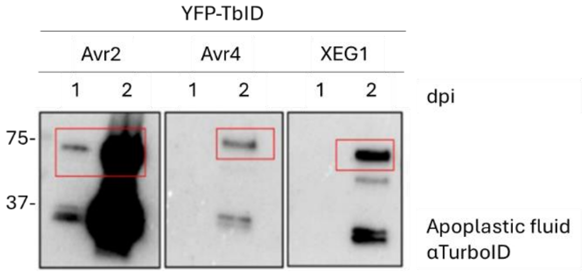


**Figure S3**. Apoplastic accumulation of effector–YFP–TurboID fusions upon their transient expression in leaves of *N. benthamiana*. Leaves of WT *N. benthamiana* plants were agroinfiltrated with Avr2, Avr4 or XEG1, all fused to YFP–TurboID (TbID), after which apoplastic fluid was collected at 1 and 2 dpi and ~11 µl of each sample was analysed by SDS–PAGE and WB, using anti-TurboID antibodies. Full-length effector–YFP–TurboID fusions, having a molecular weight of about 75kDa, were detected in the apoplast at 2 dpi for all constructs, with Avr2–YFP–TurboID already accumulating at 1 dpi and being strongly overexpressed at 2 dpi. Molecular mass markers (kDa) are indicated on the left.

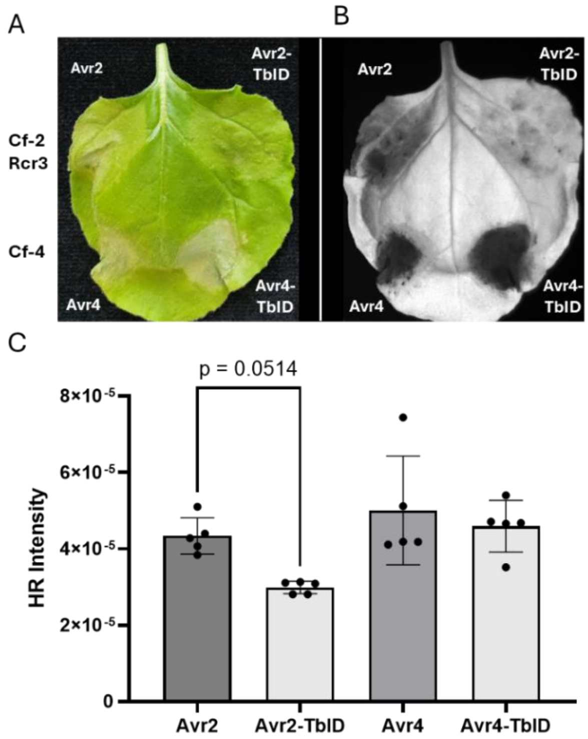


**Figure S4.** Comparison of the intensity of the HR induced by Avr2 and Avr4, with or without a TurboID fusion. (A, B) Representative HR phenotypes in *N. benthamiana* leaves transiently expressing Avr2 or Avr2–TurboID (TbID) together with Cf-2 and tomato Rcr3, or Avr4 or Avr4–TurboID together with Cf-4, as indicated. Leaves were imaged at 4 dpi under visible light and by using red epifluorescence illumination with a 695/55 nm filter. (C) Quantification of the HR intensity based on the red fluorescence signal in the infiltrated leaf areas. Each bar represents the mean ± SD of five biological replicates (infiltration spots), and individual data points are shown. Statistical significance was assessed using one-way ANOVA followed by Tukey's multiple comparison test. Note that Avr2–TurboID elicits a compromised HR when compared to Avr2 ($p$ = 0.0514), whereas no significant difference in the intensity of the HR is observed between Avr4 and Avr4–TurboID.

A

B

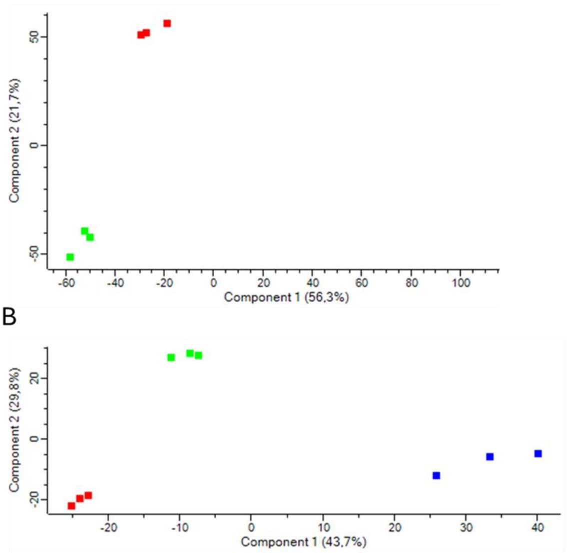


**Figure S5.** Principal component analysis of effector–TurboID PL–LC-MS/MS datasets. Principal component analysis (PCA) of label-free quantification (LFQ) intensities for all PL–MS samples. (A) Separation of Avr4-TurboID- and XEG1–TurboID-generated PL–LC-MS/MS datasets in *N. benthamiana*, based on the first two principal components. (B) PCA of the Avr2-TurboID–, Avr4-TurboID–, and XEG1–TurboID-generated PL–LC-MS/MS datasets in tomato. Biological replicates cluster tightly by bait, indicating high reproducibility, while effector-specific clusters reflect distinct proxitome compositions of Avr2, Avr4 and XEG1, which are represented with blue, green, and red squares, respectively.

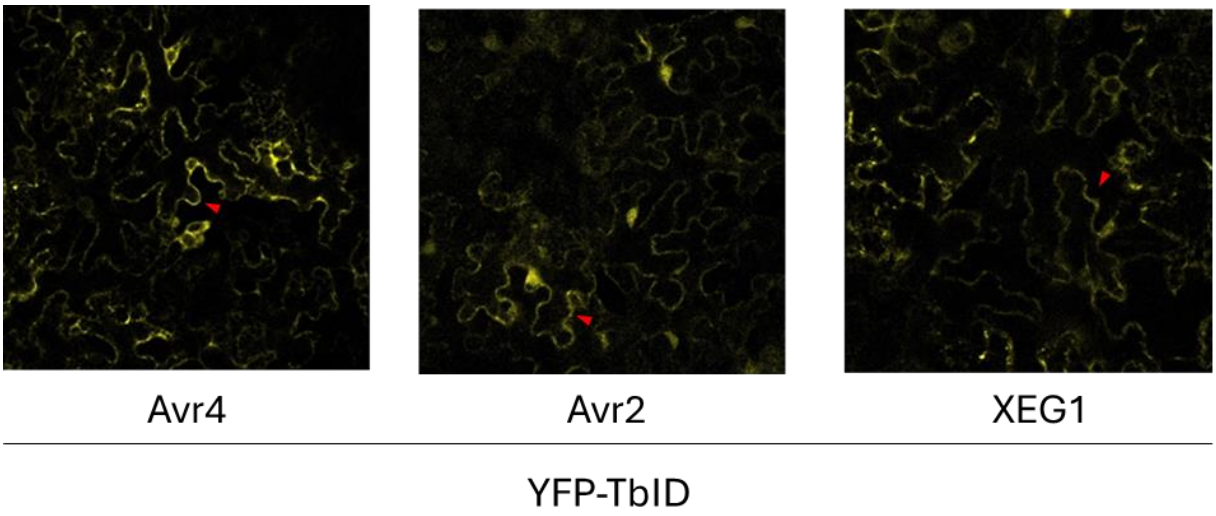


**Figure S6.** Effector–YFP–TurboID fusion proteins are properly expressed in tomato leaves. Confocal laser scanning microscopy images of tomato leaf epidermal cells expressing TurboID (TbID) fusions under control of the 35S promoter. YFP fluorescence was used to monitor protein expression and localisation by confocal microscopy. Images were taken at 2 dpi. YFP fluorescence is indicated by red arrowheads.

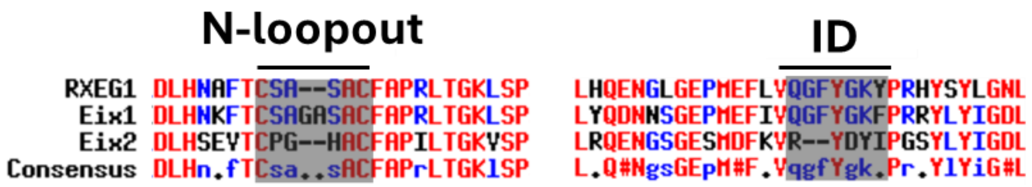


**Figure S7.** Amino acid sequence alignment of *Nb*RXEG1 and its closest tomato orthologs, *Sl*Eix1 and *Sl*Eix2. Sequence alignment of *Nb*RXEG1, *Sl*Eix1 and *Sl*Eix2 is focused on the N-loopout and ID regions. Residues previously implicated in XEG1 binding in the N-loopout region and *Nb*RXEG1 activation in the ID are indicated by grey shading. Note that the amino acid sequences of the N-loopout and the ID segments of *Sl*Eix1 closely match those of NbRXEG1, whereas *Sl*Eix2 displays various substitutions and small deletions, consistent with the structural distortions observed in AlphaFold models.

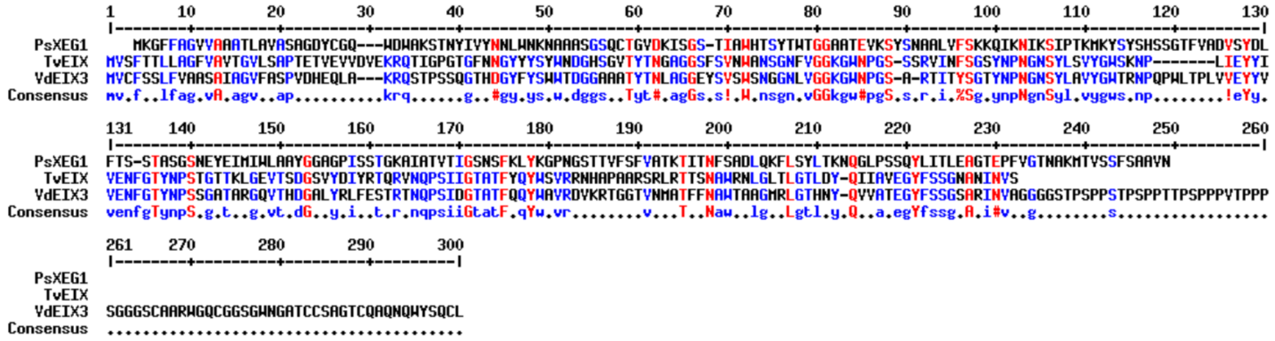


**Figure S8.** Amino acid sequence alignment of XEG1, *Tv*EIX and *Vd*EIX3. The primary amino acid sequences of the EIX elicitors have low sequence similarity with XEG1.

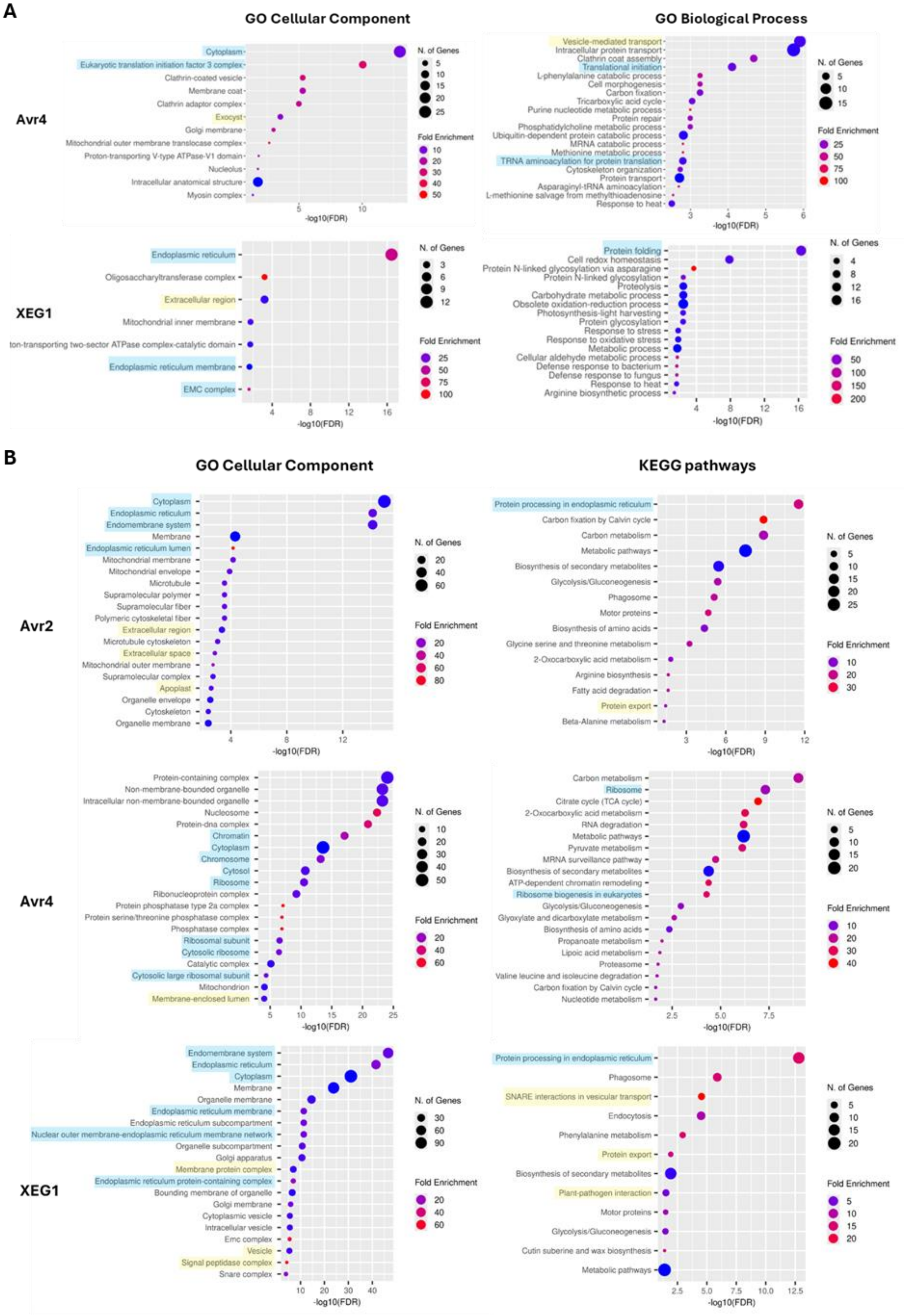
A
GO Cellular Component
GO Biological Process
Avr4
Cytoplasm
Eukaryotic translation initiation factor 3 complex
Clathrin-coated vesicle
Membrane coat
Clathrin adaptor complex
Exocyst
Golgi membrane
Mitochondrial outer membrane translocase complex
Proton-transporting V-type ATPase-V1 domain
Nucleolus
Intracellular anatomical structure
Myosin complex
Vesicle-mediated transport
Intracellular protein transport
Clathrin coat assembly
Translational initiation
L-phenylalanine catabolic process
Cell morphogenesis
Carbon fixation
Tricarboxylic acid cycle
Purine nucleotide metabolic process
Protein repair
Phosphatidylcholine metabolic process
Ubiquitin-dependent protein catabolic process
MRNA catabolic process
Methionine metabolic process
TRNA aminoacylation for protein translation
Cytoskeleton organization
Protein transport
Asparaginyl-tRNA aminoacylation
L-methionine salvage from methylthioadenosine
Response to heat
XEG1
Endoplasmic reticulum
Oligosaccharyltransferase complex
Extracellular region
Mitochondrial inner membrane
ton-transporting two-sector ATPase complex-catalytic domain
Endoplasmic reticulum membrane
EMC complex
Protein folding
Cell redox homeostasis
Protein N-linked glycosylation via asparagine
Protein N-linked glycosylation
Proteolysis
Carbohydrate metabolic process
Obsolete oxidation-reduction process
Photosynthesis-light harvesting
Protein glycosylation
Response to stress
Response to oxidative stress
Metabolic process
Cellular aldehyde metabolic process
Defense response to bacterium
Defense response to fungus
Response to heat
Arginine biosynthetic process
N. of Genes
Fold Enrichment
-log10(FDR)
B
GO Cellular Component
KEGG pathways
Avr2
Cytoplasm
Endoplasmic reticulum
Endomembrane system
Membrane
Endoplasmic reticulum lumen
Mitochondrial membrane
Mitochondrial envelope
Microtubule
Supramolecular polymer
Supramolecular fiber
Polymeric cytoskeletal fiber
Extracellular region
Microtubule cytoskeleton
Extracellular space
Mitochondrial outer membrane
Supramolecular complex
Apoplast
Organelle envelope
Cytoskeleton
Organelle membrane
Protein processing in endoplasmic reticulum
Carbon fixation by Calvin cycle
Carbon metabolism
Metabolic pathways
Biosynthesis of secondary metabolites
Glycolysis/Gluconeogenesis
Phagosome
Motor proteins
Biosynthesis of amino acids
Glycine serine and threonine metabolism
2-Oxocarboxylic acid metabolism
Arginine biosynthesis
Fatty acid degradation
Protein export
Beta-Alanine metabolism
Avr4
Protein-containing complex
Non-membrane-bounded organelle
Intracellular non-membrane-bounded organelle
Nucleosome
Protein-dna complex
Chromatin
Cytoplasm
Chromosome
Cytosol
Ribosome
Ribonucleoprotein complex
Protein phosphatase type 2a complex
Protein serine/threonine phosphatase complex
Phosphatase complex
Ribosomal subunit
Cytosolic ribosome
Catalytic complex
Cytosolic large ribosomal subunit
Mitochondrion
Membrane-enclosed lumen
Carbon metabolism
Ribosome
Citrate cycle (TCA cycle)
2-Oxocarboxylic acid metabolism
RNA degradation
Metabolic pathways
Pyruvate metabolism
MRNA surveillance pathway
Biosynthesis of secondary metabolites
ATP-dependent chromatin remodeling
Ribosome biogenesis in eukaryotes
Glycolysis/Gluconeogenesis
Glyoxylate and dicarboxylate metabolism
Biosynthesis of amino acids
Propanoate metabolism
Lipoic acid metabolism
Proteasome
Valine leucine and isoleucine degradation
Carbon fixation by Calvin cycle
Nucleotide metabolism
XEG1
Endomembrane system
Endoplasmic reticulum
Cytoplasm
Membrane
Organelle membrane
Endoplasmic reticulum membrane
Endoplasmic reticulum subcompartment
Nuclear outer membrane-endoplasmic reticulum membrane network
Organelle subcompartment
Golgi apparatus
Membrane protein complex
Endoplasmic reticulum protein-containing complex
Bounding membrane of organelle
Golgi membrane
Cytoplasmic vesicle
Intracellular vesicle
Emc complex
Vesicle
Signal peptidase complex
Snare complex
Protein processing in endoplasmic reticulum
Phagosome
SNARE interactions in vesicular transport
Endocytosis
Phenylalanine metabolism
Protein export
Biosynthesis of secondary metabolites
Plant-pathogen interaction
Motor proteins
Glycolysis/Gluconeogenesis
Cutin suberine and wax biosynthesis
Metabolic pathways
N. of Genes
Fold Enrichment
-log10(FDR)

**Figure S9.** GO and pathway enrichment of the effector–TurboID proxitomes in (A) *N. benthamiana* and (B) tomato. Gene Ontology (GO) term and KEGG pathway enrichment analyses for proteins significantly enriched by transiently expressing effector-TurboID fusions in leaves of *N. benthamiana* and tomato were performed. The dots indicate –log10 adjusted p-values for selected biological processes and cellular component terms. Enrichment of translation-related categories (blue highlighted) in the proxitome of Avr4-TurboID could reflect relatively high expression or unspecific localisation, whereas apoplast- and secretory pathway-related categories (yellow highlighted) are particularly prominent in Avr2– and XEG1–TurboID proxitomes, which could be attributed to the NbPR1a signal peptide for extracellular targeting fused to these effectors. These results suggest that replacing the native secretion signal of an effector with a plant secretion signal could increase the secretion efficiency in plants. The dot plot charts were generated by ShinyGO 0.85.1 (Ge et al., 2020), with an FDR cutoff of 0.05.

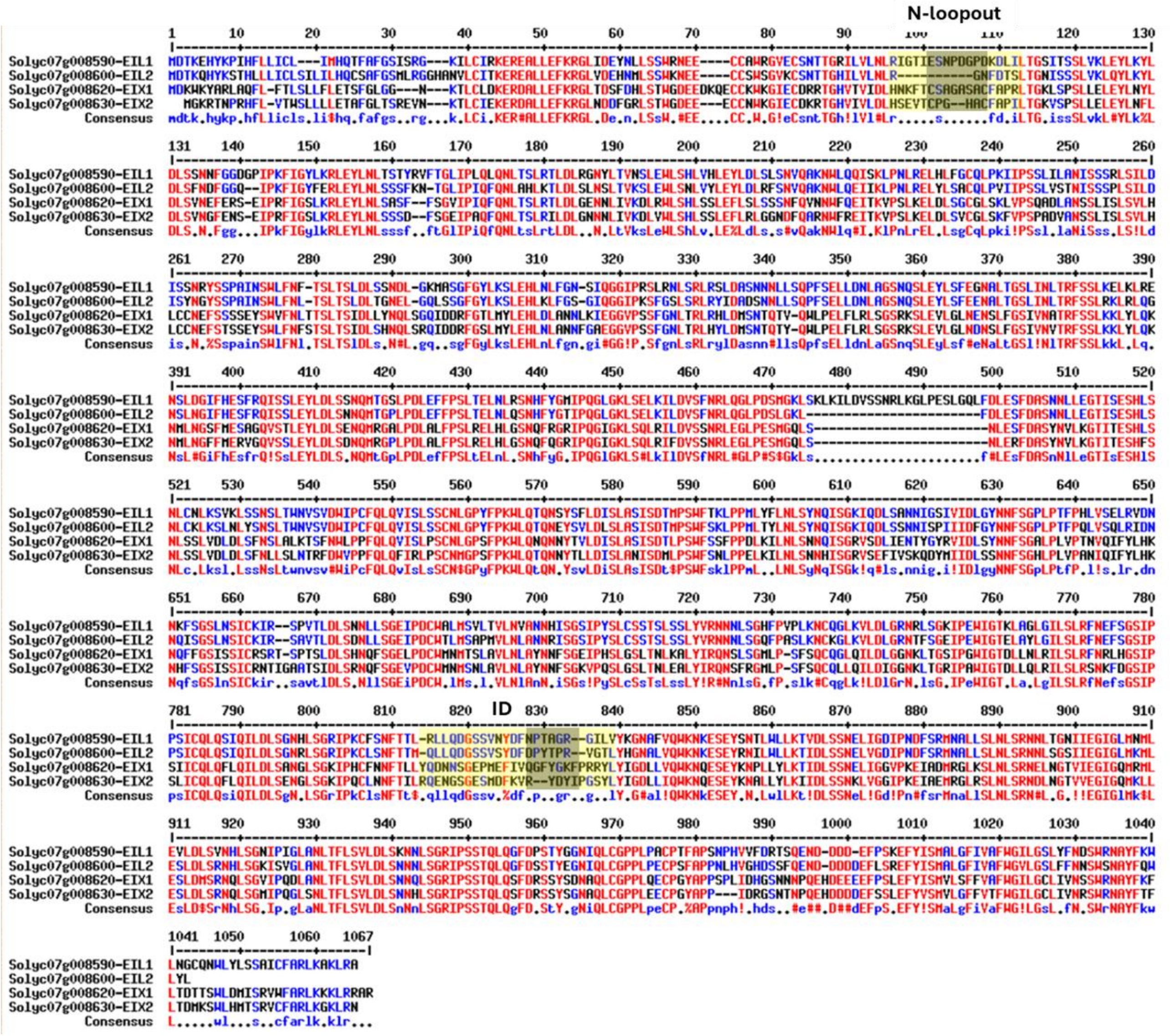

**Figure S10.** Alignment of the amino acid sequences of the RLPs encoded by the tomato *EIX* locus. Multiple sequence alignment of the RLPs that are encoded by the *EIX* locus of tomato, EIX-LIKE 1 (Eil1), Eil2, Eix1 and Eix2, with the N-loopout and ID regions highlighted in yellow. Residues earlier implicated in XEG1 binding and *Nb*RXEG1 activation are shaded in grey. Sequence variation in the effector-recognition segments indicates that the RLPs of the EIX locus may have evolved to recognise distant effectors.

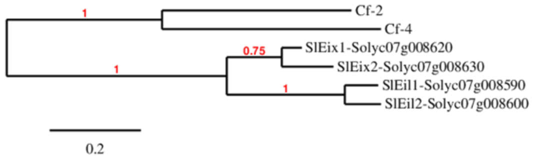


**Figure S11.** Phylogenetic relationships between the RLPs encoded by the *EIX* locus and the Cf-2 and Cf-4 proteins. The maximum-likelihood phylogeny (Dereeper et al., 2008) of tomato Eix1, Eix2, Eil1, Eil2, Cf-2, and Cf-4 is shown. The RLPs encoded by the *EIX* locus cluster together, while the Eixs and Eils branch in the cluster.

>**Cf-4**
MGCVKLVFFMLYVFLFQLVSSSSLPHLCPEDQALALLEFKNMFTVNPNASDYCYDRRTLSWNKSTSCCSWDGVHCDETTGQVIELDLRCIQLQGKFHSNSSLFQLSNLKRLDLSYN
DFTGSPISPKFGEFSDLTHLDLSHSSFRGVIPSEISHLSKLYVLRISLNELTFGPHNFELLLKNLTQLKVLDLESINISSTIPLNFSSHLTNLWLPYTELRGILPERVFHLSDLEFLDLSSNP
QLTVRFPTTKWNSSASLMKLYLYNVNIDDRIPESFSHLTSLHKLYMSRSNLSGPIPKPLWNLTNIVFLDLNNNHLEGPIPSNVSGLRNLQILWLSSNNLNGSIPSWIFSLPSLIGLDLSN
NTFSGKIQEFKSKTLSTVTLKQNKLKGPIPNSLLNQKNLQFLLLSHNNISGHISSAICNLKTLILLDLGSNNLEGTIPQCVVERNEYLSHLDLSNNRLSGTINTTFSVGNILRVISLHGN
KLTGKVPRSMINCKYLTLLDLGNNMLNDTFPNWLGYLFQLKILSLRSNKLHGPIKSSGNTNLFMGLQILDLSSNGFSGNLPERILGNLQTMKEIDESTGFPEYISDPYDIYYNYLTTIST
KGQDYDSVRILDSNMIINLSKNRFEGHIPSIIGDLVGLRTLNLSHNVLEGHIPASFQNLSVLESLDLSSNKISGEIPQQLASLTFLEVLNLSHNHLVGCIPKGKQFDSFGNTSYQGND
GLRGFPLSKLCGGEDQVTTPAELDQEEEEEDSPMISWQGVLVGYGCGLVIGLSVIYIMWSTQYPAWFSRMDLKLEHIITTKMKKHKKRY

>**Cf-2**
MMMVSRKVVSSLQFFTLFYLFTVAFASTEEATALLKWKATFKNQNNSFLASWIPSSNACKDWYGVVCFNGRVNTLNITNASVIGTLYAFPFSSLPSLENLDLSKNNIYGTIPPEIGNL
TNLVYLDLNNNQISGTIPPQIGLLAKLQIIRIFHNQLNGFIPKEIGYLRSLTKLSLGINFLSGSIPASVGNLNNLSFLYLYNNQLSGSIPEEISYLRSLTELDLSDNALNGSIPASLGNMN
NLSFLFLYGNQLSGSIPEEICYLRSLTYLDLSENALNGSIPASLGNLNNLSFLFLYGNQLSGSIPEEIGYLRSLNVLGLSENALNGSIPASLGNLKNLSRLNLVNNQLSGSIPASLGNLN
NLSMLYLYNNQLSGSIPASLGNLNNLSMLYLYNNQLSGSIPASLGNLNNLSRLYLYNNQLSGSIPEEIGYLSSLTYLDLSNNSINGFIPASFGNMSNLAFLFLYENQLASSVPEEIGYL
RSLNVLDLSENALNGSIPASFGNLNNLSRLNLVNNQLSGSIPEEIGYLRSLNVLDLSENALNGSIPASFGNLNNLSRLNLVNNQLSGSIPEEIGYLRSLNDLGLSENALNGSIPASLG
NLNNLSMLYLYNNQLSGSIPEEIGYLSSLTYLSLGNNSLNGLIPASFGNMRNLQALILNDNNLIGEIPSSVCNLTSLEVLYMPRNNLKGKVPQCLGNISNLQVLSMSSNSFSGELPSSI
SNLTSLQILDFGRNNLEGAIPQCFGNISSLEVFDMQNNKLSGTLPTNFSIGCSLISLNLHGNELEDEIPRSLDNCKKLQVLDLGDNQLNDTFPMWLGTLPELRVLRLTSNKLHGPIRS
SRAEIMFPDLRIIDLSRNAFSQDLPTSLFEHLKGMRTVDKTMEEPSYESYYDDSVVVVTKGLELEIVRILSLYTVIDLSSNKFEGHIPSVLGDLIAIRILNVSHNALQGYIPSSLGSLSIL
ESLDLSFNQLSGEIPQQLASLTFLEFLNLSHNYLQGCIPQGPQFRTFESNSYEGNDGLRGYPVSKGCGKDPVSEKNYTVSALEDQESNSEFFNDFWKAALMGYGSGLCIGISIIYILI
STGNLRWLARIIEELEHKIIMQRRKKQRGQRNYRRRNNRF

>**Eix1**
MDKWKYARLAQFLFTLSLLFLETSFGLGGNKTLCLDKERDALLEFKRGLTDSFDHLSTWGDEEDKQECCKWKGIECDRRTGHVTVIDLHNKFTCSAGASACFAPRLTGKLSPSLLE
LEYLNYLDLSVNEFERSEIPRFIGSLKRLEYLNLSASFFSGVIPIQFQNLTSLRTLDLGENNLIVKDLRWLSHLSSLEFLSLSSSNFQVNNWFQEITKVPSLKELDLSGCGLSKLVPSQA
DLANSSLISLSVLHLCCNEFSSSSEYSWVFNLTTSLTSIDLLYNQLSGQIDDRFGTLMYLEHLDLANNLKIEGGVPSSFGNLTRLRHLDMSNTQTVQWLPELFLRLSGSRKSLEVLGL
NENSLFGSIVNATRFSSLKKLYLQKNMLNGSFMESAGQVSTLEYLDLSENQMRGALPDLALFPSLRELHLGSNQFRGRIPQGIGKLSQLRILDVSSNRLEGLPESMGQLSNLESFDA
SYNVLKGTITESHLSNLSSLVDLDLSFNSLALKTSFNWLPPFQLQVISLPSCNLGPSFPKWLQNQNNYTVLDISLASISDTLPSWFSSFPPDLKILNLSNNQISGRVSDLIENTYGYRV
IDLSYNNFSGALPLVPTNVQIFYLHKNQFFGSISSICRSRTSPTSLDLSHNQFSGELPDCWMNMTSLAVLNLAYNNFSGEIPHSLGSLTNLKALYIRQNSLSGMLPSFSQCQGLQILD
LGGNKLTGSIPGWIGTDLLNLRILSLRFNRLHGSIPSIICQLQFLQILDLSANGLSGKIPHCFNNFTLLYQDNNSGEPMEFIVQGFYGKFPRRYLYIGDLLVQWKNQESEYKNPLLYLK
TIDLSSNELIGGVPKEIADMRGLKSLNLSRNELNGTVIEGIGQMRMLESLDMSRNQLSGVIPQDLANLTFLSVLDLSNNQLSGRIPSSTQLQSFDRSSYSDNAQLCGPPLQECPGYA
PPSPLIDHGSNNNPQEHDEEEEFPSLEFYISMVLSFFVAFWGILGCLIVNSSWRNAYFKFLTDTTSWLDMISRVWFARLKKKLRRAR

**Figure S12.** Peptide coverage of Cf-4, Cf-2, and Eix1 in the PL–LC-MS/MS datasets generated from tomato. The detected tryptic peptides corresponding to each receptor that was identified in the present study are highlighted in yellow. Some peptides are highlighted together because one follows the other.

**Table S1.** Number of *Sl*Eix1-derived peptides detected in each replicate of the LC-MS/MS data generated by transiently expressing the different effector-TurboID fusions in leaves of MM-Cf-2 tomato plants.

| Protein ID | Unique peptides | Avr4-TbID replicate 1 | Avr4-TbID replicate 2 | Avr4-TbID replicate 3 | Avr2-TbID replicate 1 | Avr2-TbID replicate 2 | Avr2-TbID replicate 3 | XEG1-TbID replicate 1 | XEG1-TbID replicate 2 | XEG1-TbID replicate 3 |
|---|---|---|---|---|---|---|---|---|---|---|
| Q6JN47 | 5 | †0 | 0 | 0 | 0 | 0 | 0 | 2 | 3 | 5 |

†Number of peptides matching the tomato Eix1 protein (Q6JN47), detected by LC-MS/MS analysis. Note that peptides matching *Sl*Eix1 were only identified in the replicates in which XEG1-TurboID (TbID) was transiently expressed. In all these replicates, peptides derived from SlEix1 were detected, and overall, five unique peptides were found.

**Table S2.** Nucleotide sequences of the primers used in this study.

| **Primer name** | **Sequence (5'- to 3'-)** |
|---|---|
| Primers used for linearising the TurboID plasmid | |
| pEG101-YFP-TurboID-Fw* | GTGCCTAGGGTGAGCAAGGG |
| pEG101-YFP-TurboID-Rv | GATCTCGAGCGTGTCCTCTCC |
| Primers used for generating the Avr4-YFP-TurboID construct | |
| Avr4-pEG101-YFP-TbID-Fw | †gagaggacacgctcgagatcATGCACTACACAACCCTC |
| Avr4-pEG101-YFP-TbID-Rv | cccttgctcaccctaggcacATAGCCAGGATGTCCAAC |
| Primers used for generating the PR1a-Avr2-YFP-TurboID construct | |
| Insert NtPr1a Fw _TurboID | gagaggacacgctcgagatcATGGGATTTGTTCTCTTTTCACAA |
| Insert NtPr1a Fw _TurboID | gagaggacacgctcgagatcATGGGATTTGTTCTCTTTTCACAA |
| Primers used for generating the PR1a-XEG1-YFP-TurboID construct | |
| Insert NtPr1a Fw _TurboID | gagaggacacgctcgagatcATGGGATTTGTTCTCTTTTCACAA |
| Insert NtPr1a Rv _XEG1 | cactggccgcagtagtctccACCTCCTGCACATCAACAAATTT |
| Insert XEG1 Fw_NtPr1a | ttgttgatgtgcaggaggtGGAGACTACTGCGGCCAGTG |
| Insert XEG1 Rv_TurboID | cccttgctcaccctaggcacGTTGACCGCAGCCGAGAAC |

*Fw, forward; Rv, reverse.

†Nucleotides written in lower case are complementary to the plasmid sequence, and nucleotides written in upper case anneal with the inserted gene sequence.
